# Predicting antibiotic resistance evolution by mapping the local fitness landscape

**DOI:** 10.64898/2026.11.25.769263

**Authors:** Theresa Fink, Jonas Aufdermauer, Tobias Bollenbach

## Abstract

Antibiotic resistance evolution is shaped by the distribution of fitness effects (DFE) of resistance mutations, yet measurements have been confined to small sets of mutants, single proteins, or knockout libraries. Here, we show that a map of the first mutational step toward chloramphenicol resistance in *Escherichia coli* makes subsequent evolution largely predictable. Using high-throughput barcoding, we collected more than 500 spontaneous resistant mutants and measured fitness across antibiotic concentrations. With increasing concentration, the DFE broadened and its beneficial tail changed from bounded to heavy, producing rare, exceptionally fit mutants. A dose–response scaling principle explains this pattern and predicts the fitness of each mutant at any concentration from just two mutant-specific parameters. Whole-genome sequencing linked these fitness effects to their underlying mutations, exposing a vast mutational target that is dominated by amplifications and loss-of-function mutations, reaches from efflux regulators to central metabolism, and contains variants of minimal fitness cost in every class. Population-genetic simulations built on these data, without any fitted parameters, predicted the phenotypic and mutational outcomes of longer-term evolution experiments. Mapping local fitness landscapes may thus offer a general route to forecasting antibiotic resistance evolution.

## Introduction

The discovery of antibiotics in the early 20^th^ century transformed once-deadly bacterial infections into treatable conditions^1^. However, frequent and improper antibiotic use in clinical practice and livestock farming has continuously selected for resistant bacteria^2,3^. Resistance now undermines therapy and causes deaths directly attributable to untreatable infections^3–5^. Understanding—and ideally predicting—how resistance evolves is therefore essential for preserving antibiotic efficacy and designing treatments that curb resistance evolution^6,7^.

Antibiotic resistance evolves readily: resistant lineages arise through spontaneous mutations or horizontal gene transfer of resistance genes from neighboring bacteria^8^ and then rise in frequency under antibiotic selection. Transfer of antibiotic resistance genes, such as antibiotic-degrading enzymes, can immediately confer high-level resistance. In contrast, multiple spontaneous mutations frequently need to accumulate to achieve it^9,10^. Lineage-tracking experiments using DNA barcodes have directly revealed these dynamics. Lineages carrying beneficial mutations expand and compete for fixation, a process called clonal interference^11–13^. Predicting these dynamics requires two quantities: the rate at which mutations arise and their effects on fitness in a given environment, i.e., the distribution of fitness effects (DFE). While mutation rates have been estimated in various ways and conditions^14–19^, comprehensive measurements of the DFE remain scarce.

The DFE describes the fitness effects of new mutations relative to an ancestral genotype in a given environment^20^. Because selection purges deleterious mutations, adaptation is shaped by the beneficial tail of this distribution^21,22^. Extreme value theory suggests that this tail should be exponential in well-adapted organisms^23^. Experiments partly agree^24–26^, but heavy and bounded tails have also been reported^21,27,28^. Which form is observed appears to depend on how well adapted a population is. The tail approaches an exponential during adaptation^26,29^, and deviations from it have been reported to grow with antibiotic concentration, and thus selection pressure^25^. The DFE is therefore not static but depends on both environment and genetic background^21,30^; this context dependence complicates fitness predictions and can constrain adaptive pathways^31–33^.

The 4.6-Mb *Escherichia coli* (*E. coli*) genome allows 14 million point mutations alone, before counting insertions, deletions, and amplifications. Mapping the fitness effect of each is infeasible, even ignoring gene–gene and gene–environment interactions^21^. To date, the DFE has been mapped directly only for small sets of mutants from viral or bacterial genomes or for individual proteins^25,27,34,35^. It has also been inferred indirectly from evolution experiments that track lineages with neutral markers such as DNA barcodes^11,29,36^, but there the underlying mutations usually remain unknown. Gene-knockout and transposon libraries reveal genome-wide fitness effects^26,37,38^ but are mostly restricted to loss-of-function mutations, an important but incomplete subset. Thus, despite its central role in evolutionary dynamics and in predictive models of evolution, the shape of the DFE remains poorly understood.

The rate at which resistance mutations arise is the product of the overall mutation rate and the mutational target size, i.e., the number of genetic changes that confer resistance. The mutation rate is well characterized. In *E. coli*, it is on the order of 10^−3^ per genome per generation, whereas beneficial mutation rates are two to six orders of magnitude lower^15,18,39^. Mutation rates can be elevated by loss-of-function mutations in replication-error correction or by stress-induced mutagenesis^15,17,40,41^, both repeatedly observed in antibiotic resistance evolution^40–42^. The target size, in contrast, varies across antibiotics and remains poorly characterized, as it requires identifying the mutations underlying the DFE.

Could a sufficiently complete map of the first mutational step—the local fitness landscape— make resistance evolution predictable? Here, we test this idea for resistance to the ribosome inhibitor chloramphenicol (CHL) in *E. coli*. CHL is a model antibiotic for quantitative studies of bacterial growth and resistance^9,33,43,44^. Because it is bacteriostatic, growth rate is a well-defined measure of fitness at any concentration, and resistance evolves through diverse mutations rather than a single target^9,33,37^. We isolated more than 500 spontaneous resistant mutants, which can carry any type of mutation, and combined DNA barcoding with whole-genome sequencing to link each mutant’s fitness across antibiotic concentrations to its genotype. The resulting map reveals a mutational target far larger than the set of resistance genes found in previous evolution experiments, as well as a simple scaling principle that predicts the DFE at any antibiotic concentration. Population-genetic simulations based on this map, with no fitted parameters, predict the phenotypic and mutational outcomes of longer-term evolution experiments, a key step toward forecasting resistance evolution.

## Results

### Hundreds of barcoded mutants map the first mutational step

To map the DFE for antibiotic resistance—a prerequisite for predicting resistance evolution— we developed a high-throughput approach that isolates resistant mutants, each carrying a unique DNA barcode. Starting from the barcoded ancestor library, we selected spontaneous CHL-resistant mutants at the ancestral minimal inhibitory concentration (MIC) in a Luria–Delbrück fluctuation assay^45^ (Fig. 1A and Methods). Most cultures yielded no mutant colony (Figs. 1B and 1C), so each colony most likely represents an independent mutational event. In total, we characterized 535 CHL-resistant mutants. Matching the observed distribution of colony numbers to simulations of the selection procedure (Methods) yielded a beneficial mutation rate of 1.85 × 10^−6^ per genome per generation (95 % confidence interval (CI), 1.55 × 10^−6^ to 1.93 × 10^−6^), consistent with the much less precise *p*_0_-method estimate^14^ of 8.1 × 10^−7^ (Fig. 1D). Estimates varied between independent experiments, most likely because small differences in effective CHL concentration shift which mutants are recovered: small-effect mutations are expected to be the most common^21,46^ and grow only close to the ancestral MIC, so slightly exceeding it excludes many of them. Our mutation rate estimate is therefore likely conservative, yet still lies in the upper range of prior estimates for antibiotic resistance^18^.

**Figure 1:**
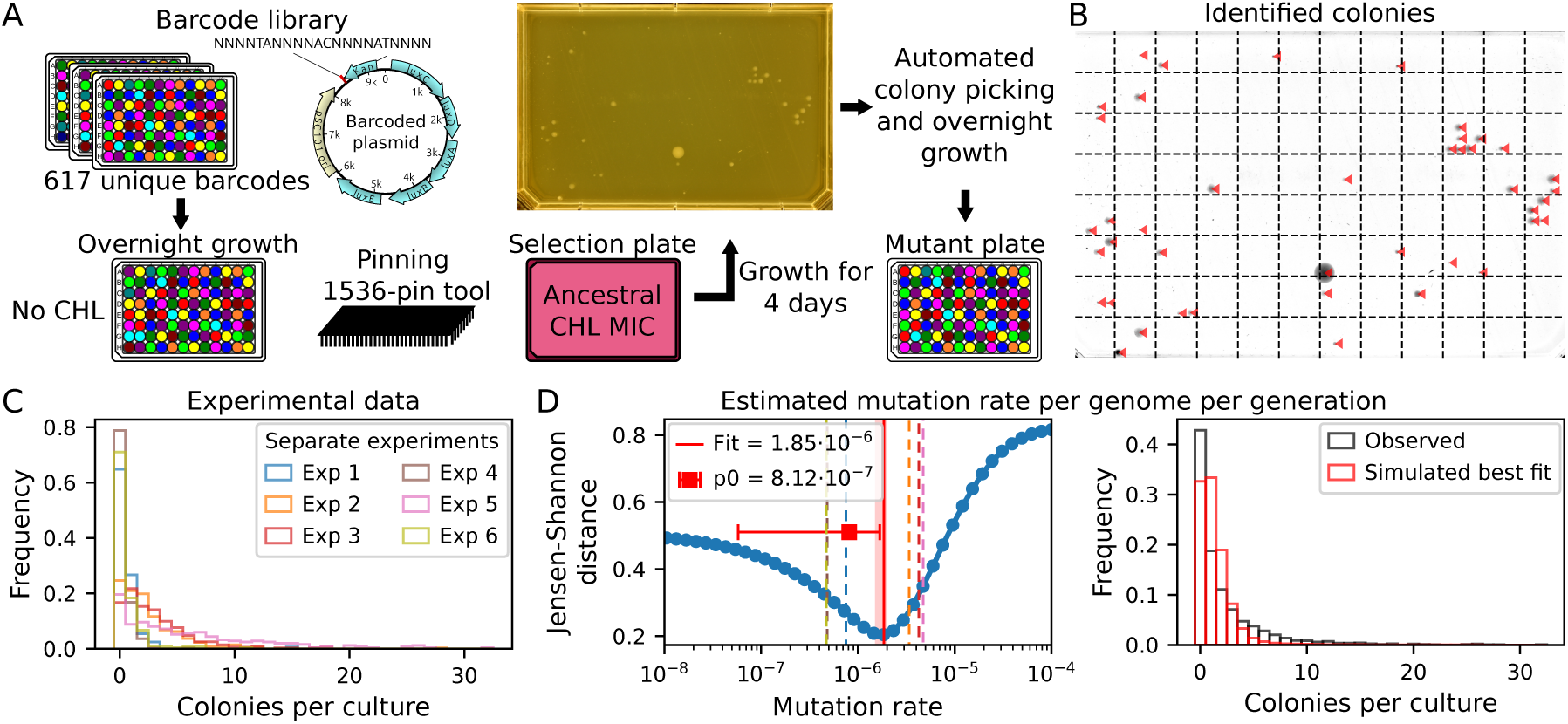
High-throughput isolation of hundreds of barcoded antibiotic-resistant mutants. **A)** Scheme of the Luria–Delbrück fluctuation assay. Independent cultures of an ancestor library with 617 unique plasmid-borne barcodes grew overnight without selection; each was then spotted twice with 16 pins onto agar containing CHL at the ancestral MIC. Colonies were collected by automated cherry picking. **B)** Example selection plate. Red arrows point to colonies identified by image segmentation. Dashed lines delimit the 16 replicate spots of each culture. **C)** Colonies per culture for each repetition of the selection experiment. **D)**Beneficial mutation rate estimated by matching experimental and simulated colony distributions (Jensen–Shannon distance). Blue line and markers, grid values of mutation rate and distance; solid red line, best fit for the full dataset; dashed lines, best fits for individual experiments; red square, *p*_0_-method estimate (*y* position arbitrary); red shading and error bars, bootstrapped 95 % CIs. Right, colonies per culture in the full dataset (black) and in simulations at the best-fit rate (red).

We used the exponential growth rate as a proxy for fitness and measured it for hundreds of mutants in pooled competition across multiple CHL concentrations, using monocultures for validation (Figs. 2A to 2C). Tracking barcode frequencies by amplicon sequencing, together with total population size, yielded each mutant’s population trajectory and thus its exponential growth rate, an absolute measure of fitness under competition (Figs. 2C and 2D and Table S2; Methods). This pooled assay mimics natural competition between the ancestor and multiple mutants. Increasing CHL concentration strongly reduced growth, as expected (Figs. 2B, 2C and S1A). In antibiotic-free medium, mutant frequencies remained nearly unchanged, indicating comparable fitness among mutants and the ancestor (Figs. 2D and S1B). At higher CHL concentrations, in contrast, a small subset of mutants rapidly rose in frequency and ultimately dominated the pool (Figs. 2D and S1B), as expected in the clonal interference regime.

**Figure 2:**
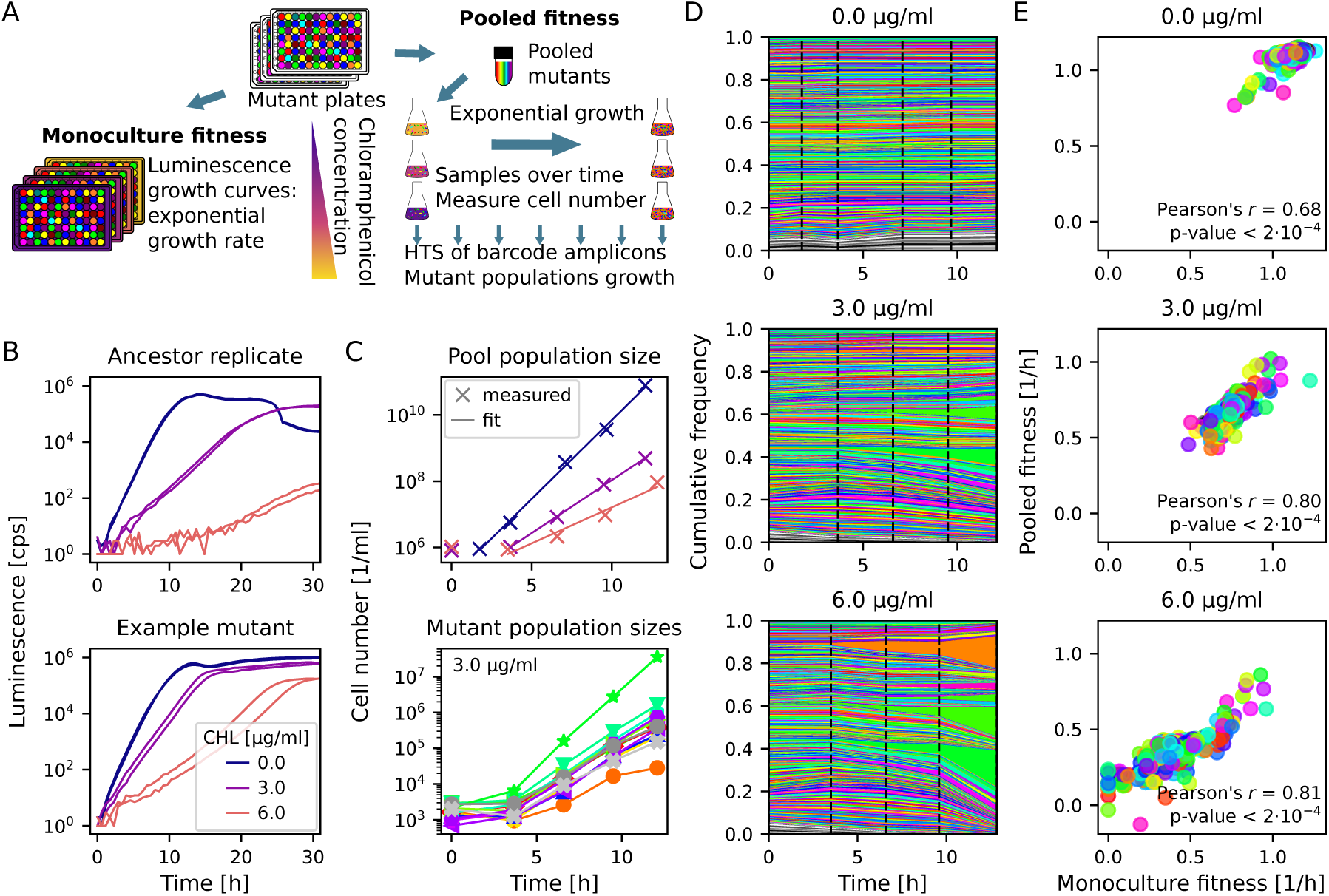
Pooled competition of barcoded mutants accurately measures fitness across antibiotic concentrations. **A)**Scheme of fitness measurements for the mutant library across antibiotic concentrations. Fitness was measured as exponential growth rate in two ways: in monoculture, from luminescence growth curves, and in pooled competition, from barcode frequencies and total cell numbers (Methods). **B)** Example luminescence growth curves of one ancestor replicate and one mutant; two technical replicates per condition. **C)** Population sizes of the pool and of individual strains over time in the competition assay, calculated from total cell numbers (top) and barcode frequencies from D (Methods); sampling time points are marked. Colored lines, twelve mutants with colors matching across panels; gray, two ancestor replicates. **D)** Muller plot of mutant frequencies (n = 474) over time at example antibiotic concentrations. Dashed lines, sampling time points; gray at the bottom, barcoded ancestor replicates (n = 18). **E)** Monoculture versus pooled-competition fitness at example antibiotic concentrations; Pearson’s *r* is given in each panel (n = 365).

We validated the pooled assay against conventional monoculture growth curves, measured by luminescence in a plate reader, and through repeated barcoding assays at a subset of concentrations (Figs. 2B, 2E and S1). Fitness values from monoculture and pooled competition correlated strongly at all CHL concentrations (Fig. 2E); the correlation was weakest without drug, where the narrow fitness range made small differences hard to resolve. This agreement likely reflects that our strain lacks CHL-degrading enzymes, so mutants interact mainly through nutrient competition and grow largely independently until nutrient exhaustion^47^. Fitness estimates were reproducible: antibiotic-free replicates within one experiment agreed closely (concordance *r*_c_ = 0.93, Fig. S1C), and separate experiments were well correlated (*r*_c_ = 0.78–0.82, Fig. S1D). The barcode-based competition assay thus precisely maps the fitness of hundreds of mutants at finely resolved antibiotic concentrations.

### Antibiotic stress broadens the distribution of fitness effects

With increasing CHL concentration, fitness differences between mutants and ancestor replicates grew more pronounced, while the median fitness decreased, as expected (Fig. 3A). At the lowest concentrations, the ancestor was slightly fitter than most mutants, but its rank dropped at higher concentrations and then remained low (Figs. 3A and 3B). Many mutants that were highly fit without antibiotic declined similarly, while others climbed in rank. Some ranked high at all concentrations, combining resistance with little or no fitness cost (Fig. 3B). These rank changes confirm that many resistance mutations are mildly deleterious without antibiotic yet beneficial in its presence^48–51^, so which mutants win depends on drug concentration.

**Figure 3:**
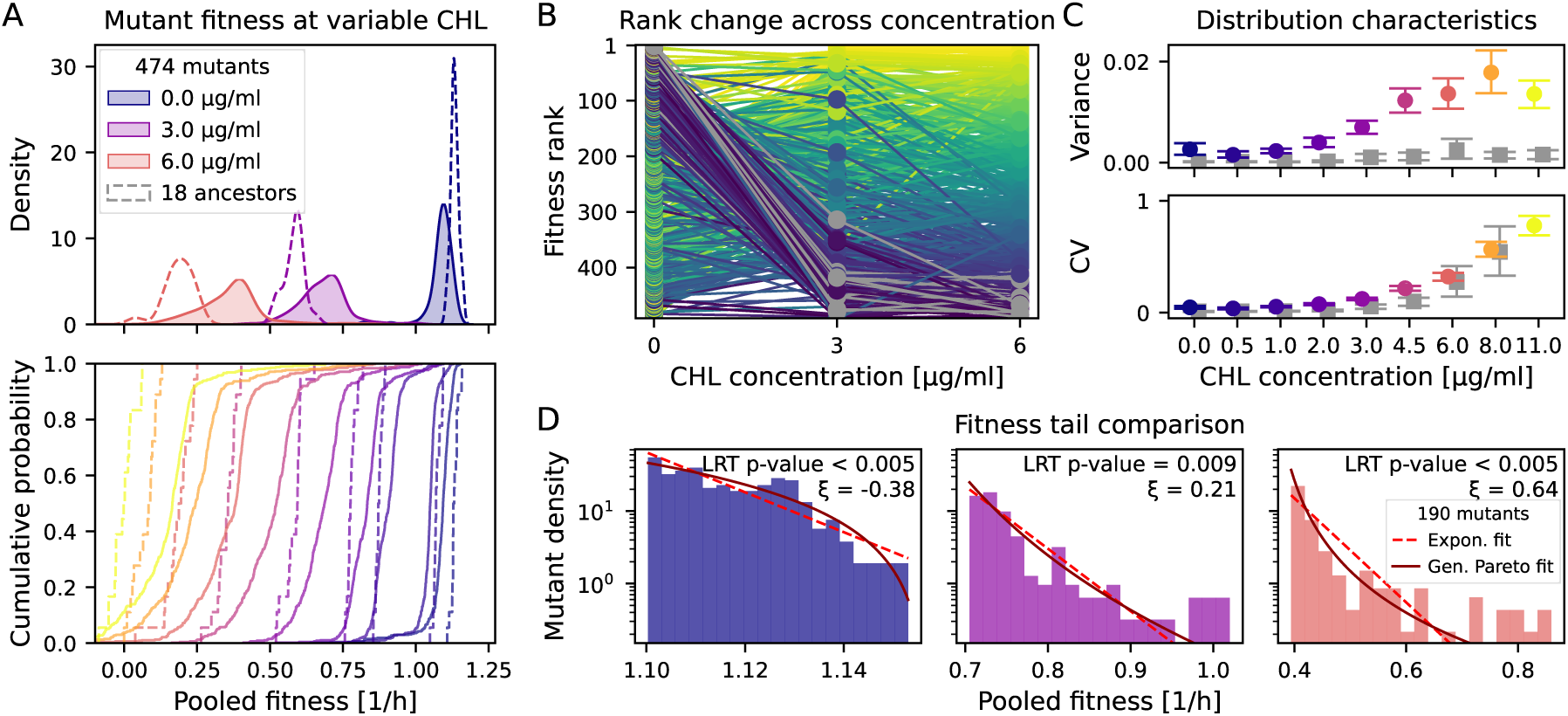
Antibiotic stress broadens the distribution of fitness effects and creates a heavy beneficial tail. **A)** Distributions of competitive fitness of 474 mutants and 18 ancestor replicates across concentrations, shown as kernel density estimates for example concentrations and empirical cumulative distributions for all. **B)** Pooled fitness rank of mutants and ancestor replicates across concentrations (rank one is the fittest at each concentration); colored by rank at 6 µg ml^−1^; gray, ancestor replicates. **C)** Variance and coefficient of variation (CV) of fitness among mutants and among ancestor replicates (gray) across all concentrations. Error bars, bootstrapped 95 % CIs. At the highest concentration, the mean ancestor fitness is negative, so its CV is not meaningful. **D)** Beneficial tail of the distribution (histogram, fittest 40 % of mutants) at example concentrations compared with a fitted exponential (dashed red line) and generalized Pareto (solid dark red line). *p*-values from a likelihood-ratio test against the exponential^46^. *ξ*, shape parameter from the generalized Pareto fit. Full data in Fig. S2.

The DFE broadened markedly with increasing CHL concentration, in both absolute and relative terms: its variance rose up to 6.8-fold and its coefficient of variation up to 16.6-fold relative to the no-drug condition (Figs. 3A and 3C). Although the deleterious tail of the DFE is not observable here, because deleterious mutants rarely form detectable colonies during isolation (Methods), this broadening aligns with the view that stress amplifies the phenotypic effects of genetic variation^52,53^.

The beneficial tail of the DFE contains the high-fitness mutations that drive adaptation; extreme-value theory suggests it to be exponential in well-adapted organisms^23^. We tested this prediction by fitting a generalized Pareto distribution to the tail^46^. Without antibiotic, the tail was bounded, with fewer beneficial mutations than expected (Fig. 3D), although our selection for resistance may have excluded mutations that are beneficial only without drug. With antibiotic above 0.5 µg ml^−1^, the exponential fit consistently underestimated the abundance of highly beneficial mutations, and a positive shape parameter indicated a heavy tail in the Fréchet domain (Figs. 3D and S2A). Across all thresholds, *ξ* increased significantly with concentration; the correlation was weakest at the 0.6 quantile used throughout (Spearman’s *r*_s_ = 0.72, *p* = 0.042, *n* = 9; Fig. S2B). Antibiotic stress thus shifts the tail from a deficit to an excess of high-fitness mutations, consistent with rare but potent resistance mutations. This shift matches earlier observations: the tail is close to exponential in populations well adapted to their environment^25,26,29^ but heavy under a stress to which they are poorly adapted and have low fitness, as for TEM-1 *β*-lactamase mutations under cefotaxime^27,54^.

### A scaling principle predicts fitness at any antibiotic concentration

Beyond mapping DFEs, the precise fitness measurements from our barcoding assay yielded dose–response curves (DRCs) of individual mutants at high resolution and throughput. We measured 492 DRCs, including ancestor replicates, across nine CHL concentrations spanning the range where the antibiotic increasingly inhibits growth (Fig. 4A and Table S2). Resistance levels varied widely and clearly exceeded those of the ancestor; the most resistant mutants were barely inhibited even at the highest concentration (Fig. 4B). To condense these measurements into a few parameters, we fitted a Hill function to the growth rate *g* as a function of concentration *c*,

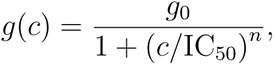

where *g*_0_ is the antibiotic-free growth rate, the half-maximal inhibitory concentration (IC_50_) is the resistance level, and *n* is the Hill exponent, which sets the steepness of the DRC, i.e., the sensitivity of growth to increasing antibiotic (Fig. 4B). This reduction to three parameters per mutant enabled us to systematically analyze relationships among key features of the DRCs.

**Figure 4:**
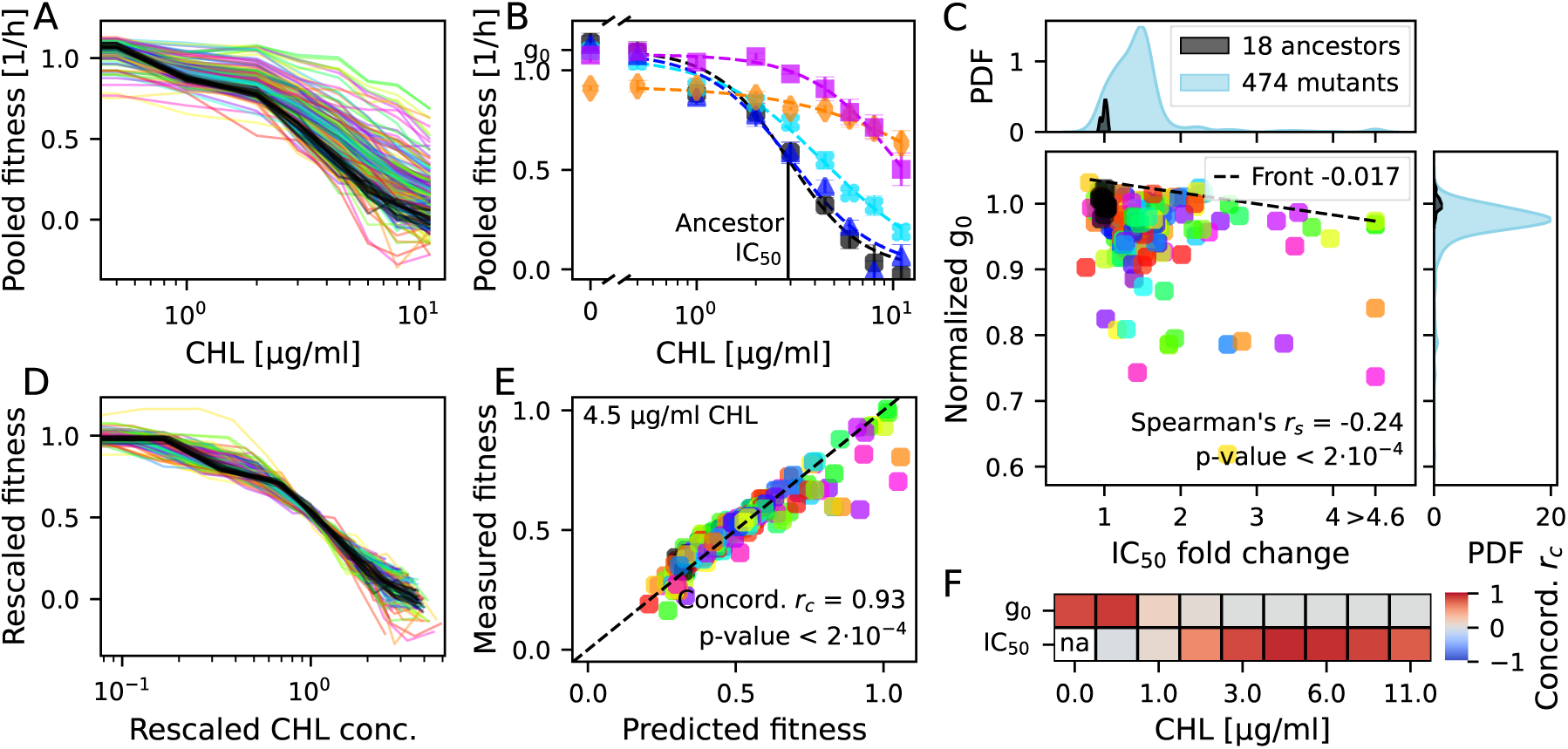
A scaling principle for dose–response curves predicts mutant fitness from two parameters. **A)** Dose-response curves of all 474 mutants (colors) and 18 ancestor replicates (black). **B)** Example dose–response curves of four mutants (colors) and one ancestor replicate (black). Dashed lines, Hill function fits; error bars, standard error of the fitted growth rate (Methods). The ancestor’s antibiotic-free fitness (*g*_0_) and IC_50_ are marked on the *y* and *x* axes, respectively. **C)** Mutant *g*_0_ versus IC_50_, both relative to the ancestor mean. Black line, linear slope of the Pareto front for low cost mutations. Fold change *>* 4.6 could not reliably be determined given our maximum, measured concentration. Top and right, kernel density estimates of the marginal distributions. **D)** Dose-response curves from A, with fitness rescaled by each strain’s *g*_0_ and CHL concentration by its IC_50_. **E)** Measured versus predicted fitness at 4.5 µg ml^−1^ CHL. Predictions use each mutant’s IC_50_ with the ancestor’s Hill exponent *n* and *g*_0_ in the Hill function (Methods). **F)** Concordance correlation coefficients (*r*_c_) between measured and predicted fitness across strains for each concentration; predictions use either the mutant’s *g*_0_ or its IC_50_, and ancestor values for the other parameters. Predicting 0 µg ml^−1^ with IC_50_ is not applicable (na).

Although some resistant mutants incurred a fitness cost, the global trade-off between resistance and antibiotic-free fitness was weak. Across all mutants, resistance (IC_50_) rose by a median factor of 1.42 (95 % CI, 1.39–1.45), while antibiotic-free fitness (*g*_0_) decreased by 2.4 % (2.2–2.6 %) compared to the ancestor (Fig. 4C). A subset of mutants had substantially lower antibiotic-free fitness or higher resistance, and a few exceeded the maximum detectable resistance of 13.5 µg ml^−1^ (Fig. 4C). Still, resistance levels generally remained below those reached when multiple mutations fix in longer evolution experiments^33,37^. Resistance and antibiotic-free fitness were weakly anticorrelated (Spearman’s *r*_s_ = 0.24, *p <* 2 10*^−^*^4^, permutation test; Fig. 4C), and many mutants achieved high resistance while retaining near-ancestral antibiotic-free fitness. Along the Pareto front, *g*_0_ fell by only 1.7 % per twofold increase in resistance (Fig. 4C).

Like antibiotic-free fitness, the steepness of the antibiotic response was largely unaffected by resistance mutations. Although DRC steepness varies widely across antibiotics^37^, most gene knockouts and the smaller sets of other mutations examined leave it unchanged for a given antibiotic^37,51,55^. To test whether this also holds for our resistant mutants, we rescaled each mutant’s DRC by its IC_50_ and *g*_0_. The rescaled curves largely collapsed onto the ancestor’s DRC (Fig. 4D). On average, the Hill exponent *n* was lower for mutants than for the ancestor, but this difference vanished when the fit was restricted to the lower inhibition levels the mutants actually reached (below 80 % inhibition; Fig. S3). Thus, DRCs of resistant mutants obey a simple scaling principle that links them to the ancestral DRC. Two parameters, IC_50_ and *g*_0_, suffice to predict a mutant’s fitness at any drug concentration. Because these two parameters fix the whole DRC, they also fix the DFE. Fitness predicted from *g*_0_ and IC_50_ alone reproduces the shift of the beneficial tail from bounded to heavy (*ξ* from −0.27 without drug to 0.70 at 11 µg ml^−1^; Fig. S4), matching the measured *ξ* with *r*_c_ = 0.78, *p* = 0.008.

Depending on the antibiotic concentration, even a single parameter, antibiotic-free fitness (*g*_0_) or resistance (IC_50_), was sufficient to describe fitness. For instance, fitness at 4.5 µg ml^−1^ CHL, predicted from the rescaled DRC using only the mutant-specific IC_50_ and the ancestor’s *g*_0_ and *n*, correlated strongly with measured fitness (Fig. 4E; Methods); larger deviations arose only for highly resistant mutants with exceptionally high fitness costs. In contrast, fitness at low drug concentrations was governed primarily by *g*_0_ (Fig. 4F). The transition between the two regimes occurred at around 1 µg ml^−1^, the concentration at which the ancestor’s growth began to decline (Fig. 4F). The fitness of resistant mutants under antibiotic stress can thus be predicted from a handful of readily accessible measurements, which we will use below to parameterize simulations of resistance evolution.

### Amplifications dominate a vast mutational target

To map the genome-wide mutational target of CHL resistance and dissect the role of different mutation types, we sequenced the genomes of 434 mutants and 15 ancestor replicates, identifying single-nucleotide polymorphisms (SNPs), small insertions and deletions, mobile-element insertions, large deletions, and amplifications (Figs. 5A and 5B, Methods). Unlike high-resolution lineage tracking^11,12^, this approach links the phenotype of each barcoded mutant to its underlying mutations (Tables S1 and S2). Most mutants (60 %) carried a single mutation (Fig. 5A), as expected for a selection designed to capture first-step mutants. A further 38 mutants (9 %) harbored two or more mutations, some of which may have arisen during growth on selection plates (Figs. 1A and 5A). Their phenotypes did not differ significantly from those of single mutants (Fig. S5A). Mutations other than amplifications were mostly present at near-complete frequency within each sequenced clone, with no significant relationship between frequency and phenotype (Fig. S6). In the remaining 137 clones, we detected no mutation (Fig. 5A), despite their increased resistance. Although some may be ancestral cells that survived selection (Methods), the IC_50_ of these isolates was significantly higher than that of ancestor replicates, though lower than that of other mutants (Figs. 5C and S5A), indicating that many carry undetected mutations. We hypothesize that many are amplifications lost during antibiotic-free growth before sequencing (Methods), as amplifications revert readily^56^. Indeed, the amplifications we did detect had a median copy number of only 1.46, indicating that many cells had already lost them—consistent with the lower fitness we observed at higher copy number (Fig. S6). Thus, the first step in resistance evolution may often involve unstable, reversible genetic changes.

**Figure 5:**
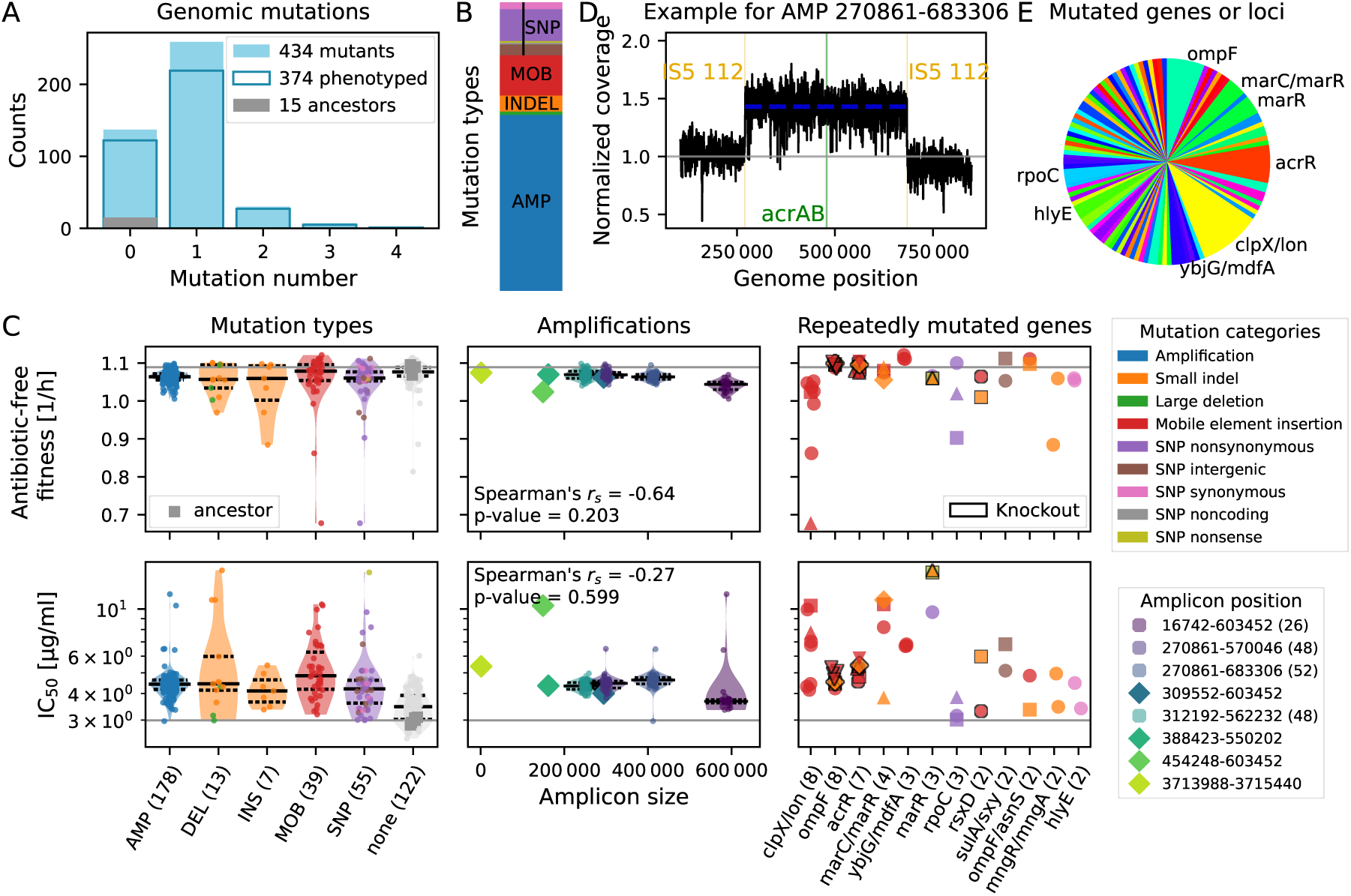
Whole-genome sequencing reveals a large and diverse mutational target dominated by amplifications. **A)** Number of mutations per sequenced mutant, and number of these mutants with phenotype data. **B)** Frequency of mutation types. **C)** Antibiotic-free fitness (*g*_0_) and resistance (IC_50_), from Hill function fits, by mutation type (left), for amplifications (middle), and for repeatedly mutated genes (right) for all phenotyped mutants including mutants with multiple mutations. Lines in violin plots, median (solid) and quartiles (dashed); gray line, ancestor median. Middle, the eight unique amplifications, sorted by size; violin plots are shown for the four that occurred repeatedly. The Spearman correlation between amplicon size and *g*_0_ or IC_50_ was computed on the medians of unique amplicons, weighted by the square root of the number of mutants carrying each; *p*-values are from a permutation test. Right, individual mutants; marker shapes distinguish mutations, and black edges mark knockouts predicted by breseq. **D)** Read coverage of an example amplified region, normalized to the median genome coverage (mutant lp5_E11_CHL_20231024_1; Table S1). Blue dashed line, median copy number of the amplicon; orange lines, amplicon borders (flanked by two copies of the same insertion sequence (IS) element); green lines, the resistance genes *acrA* and *acrB*. **E)**Mutated genes and intergenic regions, excluding amplifications and large deletions, which affect multiple genes. Genes mutated more than twice are labeled.

Consistent with this view, amplifications were the dominant cause of resistance, in line with their proposed general role in antibiotic resistance^57^. They accounted for 61 % of all detected mutations, followed by SNPs at 18 % and mobile elements (MOBs) at 14 %; insertions and deletions were rare, at 7 % (Fig. 5B). The share of amplifications is likely an underestimate, since many clones without a detectable mutation may have lost one. Most amplifications clustered in a hotspot between genomic positions 454 000 to 550 000, flanked by existing or newly inserted IS elements (Figs. 5D and S7A). This implicates genes in this region as key targets, such as the known CHL resistance genes *acrA* and *acrB*^58^, which encode components of the AcrAB-TolC multidrug efflux pump (Figs. 5D, S7A and S7C). Mutation type had only a limited effect on resistance (IC_50_) and antibiotic-free fitness (*g*_0_); median changes were small (Fig. 5C). Significant differences between single mutants were restricted to two pairwise comparisons for IC_50_ and one for *g*_0_ (Fig. S5A). Since mutation types differ little in their effects, their frequencies likely reflect intrinsic differences in mutation rates. Combined with the measured beneficial mutation rate, these frequencies give mutation rates of 1.1 × 10^−6^ for amplifications, 3.4 × 10^−7^ for SNPs, 2.6 × 10^−7^ for MOBs, and 1.0 × 10^−7^ for indels per genome per generation.

The mutated genes were diverse and widely distributed across the genome (Fig. 5E and Fig. S7A). Excluding large deletions and amplifications that hit multiple genes at once, we identified 130 mutations across all mutants, 108 of them unique, in 84 genes and intergenic regions (Fig. 5E; Table S1; Methods). Sixteen of these loci were hit more than once (Figs. 5E and S7B), often by distinct mutation types or at distinct sites, although identical SNPs (in *glnH*, *hlyE*, and *yejE*) and mobile-element insertions at identical positions also recurred. Loci hit repeatedly conferred higher resistance than loci hit once (Mann–Whitney *U* test, *p* = 3 10*^−^*^4^) but did not differ in antibiotic-free fitness (*p* = 0.74; Figs. 5C and S5B). This likely reflects a mild bias of our isolation procedure, as more resistant mutants grow more reliably into pickable colonies (Methods). Knockouts accounted for 26 % of mutations and arose through several mutation types, highlighting the many routes to loss of gene function (Figs. 5C and S7B). This fraction excludes promoter disruptions and is therefore likely an underestimate, since 35 % of mutations fell in intergenic regions, and 80 % of these were classified as promoter mutations, which can raise or lower gene expression depending on context^59^. Our approach thus revealed a wide range of genes and mutational mechanisms underlying spontaneous CHL resistance, many of which are missed by approaches restricted to loss-of-function mutations or single proteins. Much of this variation stays hidden in longer evolution experiments, as we show below.

Most repeatedly mutated genes were associated with drug efflux, in particular with the AcrAB-TolC pump, a central resistance hub^60^. Although we recovered only a single SNP in *acrA*, many mutations plausibly increase *acrAB* expression: loss-of-function mutations in the repressors *marR* and *acrR*, mobile-element insertions in the *marR* promoter (Figs. 5E and S7B), and amplification of the hotspot containing *acrA* and *acrB*. Further mutations may act indirectly: promoter mutations in *clpX* /*lon* may lower levels of the protease Lon, which degrades the activator MarA^61^. Four independent mutations hit the *rsx* operon, encoding the SoxR reduction complex, whose loss causes constitutive expression of SoxS, a second activator^62,63^. Beyond AcrAB-TolC, we found mutations in *mdfA*, a second multidrug efflux pump whose overexpression confers CHL resistance^64^, and in *glnH* and *yejE*, ATP-binding cassette (ABC) transporter subunits with unclear mechanism but links to resistance to other antibiotics^65,66^. Effects for efflux mutations differed. Loss-of-function mutations in *acrR* and promoter mutations in *mdfA* conferred intermediate resistance at negligible cost, whereas *marR* mutations conferred high resistance at a slight fitness cost (Fig. 5C). Efflux is thus a major route to CHL resistance, reached through diverse genes and mutation types—from canonical regulators to previously unrecognized transporters—and sampled often enough to estimate how frequently each arises (Fig. S7B).

Beyond efflux, many resistance-associated genes had no established link to antibiotics. Most unexpectedly, recurrent synonymous SNPs in *hlyE*, encoding the pore-forming virulence factor hemolysin E^67^, slightly increased resistance (Fig. 5C). Small indels in the promoter region shared by the sugar transporter *mngA* and its repressor *mngR* drove moderate resistance gains with highly variable fitness costs across replicates (Fig. 5C); such mutations have been observed under kanamycin selection^68^, but their role remains unclear. Metabolic genes were also well represented, including central carbon metabolism (*aceA*, *acs*, *eda*, *yggF*, *ppsR*) and the ATP synthase operon, which was hit four times independently (*atpA*, *atpD*, *atpH*, and the *atpI* promoter), echoing recent reports that mutations in core metabolic genes confer clinically relevant resistance^66,69,70^.

Among the expected targets, loss-of-function mutations in the porin *ompF* (Fig. 5E) conferred intermediate resistance without detectable cost (Fig. 5C), consistent with reduced antibiotic influx across the outer membrane^60^. As with efflux, porin function was hit at several levels: *ompF* itself and its promoter, its two-component regulator *ompR*, and a large deletion extending into *ompC*. Chloramphenicol target mutations were rare. One clone carried a point mutation in domain V of the 23S rRNA (*rrlD*), which forms the CHL binding site, and another a mutation in the 5S rRNA (*rrfE*). With seven rRNA operons in *E. coli*, a mutation in a single copy can confer only partial resistance. In contrast, nonsynonymous SNPs in the RNA polymerase subunit *rpoC*, common in evolution experiments in rich media^71^, imposed a fitness cost with minimal resistance gain (Fig. 5C), but *rpoC* mutations were also found when evolving *E. coli* to CHL with glucose, indicating a nutrient dependence^72^. These examples, together with many further mutations (Table S1), reveal a mutational target for resistance far larger and more diverse than the canonical resistance genes.

### Longer-term evolution uses few of the available mutations

Do these diverse first-step mutations also appear in populations evolved over longer time-scales? We compared our mutants with 47 *E. coli* populations carrying various gene deletions that were evolved under CHL for 180 h using a feedback-controlled protocol that maintains strong selection and tightly controls population size^33^. If clonal interference erodes the initial diversity, only the strongest mutations should fix. Indeed, the evolved populations carried together mutations in only 44 genes and intergenic regions (from 164 mutations), compared with 84 from 130 in our library (Figs. 5E, 6A and S8A). About two thirds of these 164 mutations were also found at the gene level in our library, concentrated in just 8 of the 44 loci and including the most frequently hit genes *marR*, *acrR*, and *ybjG*/*mdfA*, often by similar mutation types (Figs. 6A, S8B and S9A). Mutations in these shared genes conferred significantly higher resistance levels than mutations in genes absent from the evolved populations but did not differ in fitness cost (Fig. S9B). Thus, only a small fraction of the vast initial reservoir is fixed during longer evolution under antibiotic selection: the mutations that confer the greatest resistance.

**Figure 6:**
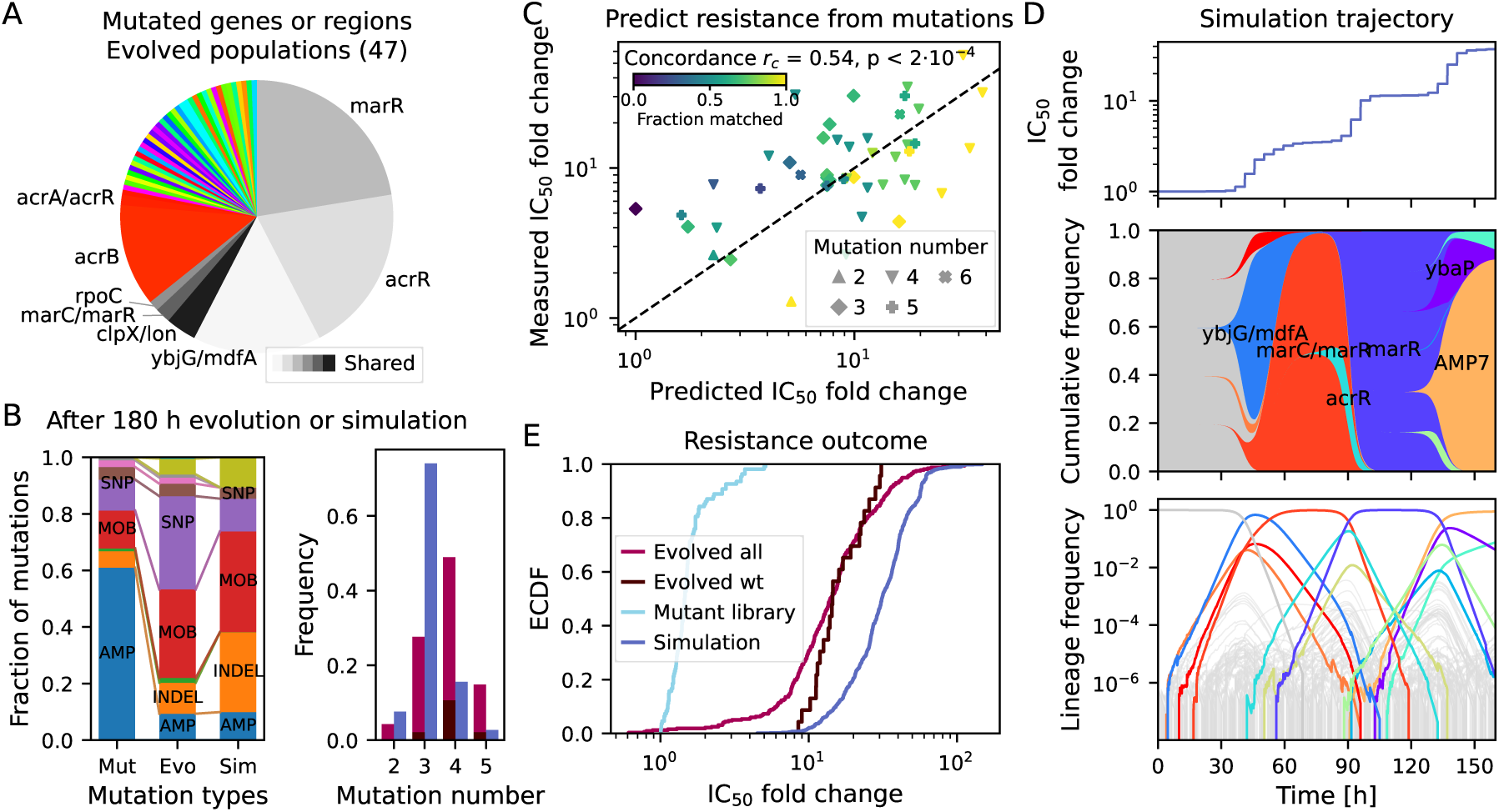
First-step mutants enable parameter-free predictions of longer-term evolutionary outcomes. **A)** Genes and intergenic regions mutated in populations sequenced after 180 h of evolution under CHL^33^. Gray shading: genes also hit in the mutant library. **B)** Left: frequencies of mutation types in the mutant library, evolved populations, and the simulations; lines connect segments of the same type. Right: number of mutations per population after 130 generations in evolved populations and simulations. Colors as in E. **C)** Measured versus predicted IC_50_ fold change of evolved populations. Each prediction multiplicatively combines the IC_50_ fold changes of the mutations in that population weighted by their frequency. Mutation IC_50_ fold changes were determined from the mutant library; mutations absent from the library were omitted (Methods). Measured IC_50_ fold changes for evolved populations are from the previous study. Agreement was assessed with the concordance correlation coefficient (*r*_c_) and a permutation test. **D)** Example simulation run. Top, adaptive increase in population resistance (IC_50_ fold change; Methods); middle, Muller plot of lineages above 0.5 % frequency, with new mutations labeled for several lineages (AMP7 is AMP388423-550202); bottom, frequencies of all lineages, with colors as in the Muller plot. Gray lines show lineages lost without reaching appreciable frequency. **E)** Cumulative distributions of IC_50_ fold change in the mutant library, all evolved gene deletion populations, only the evolved wild-type populations (Δ*lacA*), and the simulations.

Amplifications were less common in the evolved populations (11 % of all mutations; Fig. 6B). Some matched the *acrAB* region amplified in our library, but many covered a smaller region around *mdfA*, either without flanking IS elements or with newly inserted ones, and often reached higher copy numbers (Figs. S8A and S8C). Longer selection thus appears to favor more stable and targeted amplifications, as previously reported for *ampC*^73^.

Many of the available first-step resistance mutations were never selected, even as evolution proceeded and multiple resistance mutations accumulated. For example, eight first-step mutants carried *ompF* knockout or promoter mutations, which came at no detectable fitness cost but conferred only intermediate resistance, and none fixed in the evolution experiment. Epistasis could explain this, if earlier resistance mutations diminish the gain provided by *ompF*, but *ompF* mutations were also rare in our simulations, which include no epistasis (see below). Conversely, about one third of the mutations in the evolved populations, such as those in *acrB*, never appeared in our first-step mutant library (Fig. 6A). These may become beneficial only after other mutations have occurred; for example, *acrB* mutations may require increased expression of the AcrAB-TolC pump. Indeed, in evolved populations *acrB* mutations always occurred together with at least one mutation or gene deletion likely increasing *acrB* expression (e.g., knockouts of *acrR* or *marR*). Alternatively, they may simply be rare: the *acrB* mutations comprise several distinct non-synonymous SNPs, a mutation type far less common in our library than amplifications (Figs. 5B and 6B).

### Parameter-free simulations predict longer-term evolution

Resistance effects measured in our first-step library predicted the resistance gained by the evolved populations. We matched the mutations in the evolved populations with those in the library, then multiplied the resistance gains from each individual mutation to predict each population’s fold change in resistance. These predictions agreed reasonably well with the measured fold changes from the evolution experiment (concordance correlation coefficient *r*_c_ = 0.54, *p <* 2 10*^−^*^4^; Fig. 6C), even though often only a subset of the mutations could be matched. In these cases, the prediction should underestimate the resistance gain. Notably, if all mutations in a population were matched, predictions tended to exceed the observed resistance level, possibly reflecting global diminishing-returns epistasis^32,74^. In partially matched populations, these two opposing effects may partly offset each other. Thus, single-mutant measurements capture a substantial part of the resistance gained during longer evolution, even without accounting for epistasis.

To test whether the first-step mutant library can predict evolutionary trajectories, we reproduced the evolution experiment in Monte Carlo simulations (Methods). The simulations combine the measured mutation rate and DFE (Figs. 1 and 3), the DRC scaling principle (Fig. 4), and the characterized resistance mutations (Fig. 5). Simulated populations consist of lineages that grow, mutate, and are diluted, while the antibiotic concentration is adjusted to maintain 50 % growth inhibition (Fig. 6D top), mimicking the feedback control of the evolution experiment^33^. Each mutation is drawn from the library and founds a new lineage; following Das *et al.*^51^, the IC_50_ and *g*_0_ fold changes of successive mutations are multiplied without epistatic interactions in these parameters, so that predictions rest on the first-step landscape alone. Fitness is nevertheless epistatic, because it depends nonlinearly on these two parameters through the Hill function. All parameters describing mutations—their rate and their effects on fitness and resistance—are fixed by the library; none were fitted to the outcome of the evolution experiment. In the simulations, most new mutations are lost stochastically, and of the lineages that establish, only a few reach appreciable frequency (Fig. 6D), as expected under clonal interference.

Our simulations predicted the increase in resistance as well as the number, types, and target genes of the mutations that fix in longer evolution experiments. Across 1000 simulation runs, typically two to five mutations exceeded the 30 % frequency threshold of the sequencing analysis (Methods), as in the evolved populations, although the median was slightly lower (three versus four; Fig. 6B). The population structure at the time of sequencing matched as well. In both evolved and simulated populations, most detected mutations had fixed, and the rest formed a tail of intermediate-frequency mutations carried by subpopulations (Fig. S10A). The simulations thus capture not only which mutations win, but also how far their competition has progressed by the time of sampling. The predicted resistance gain was within about a factor of two of the observed one (Fig. 6E). This overestimate likely reflects diminishing-returns epistasis, which reduces the resistance gain when mutations are combined and which the simulations do not include. This agreement is notable because the predictions are absolute: no parameter was fitted to the results of the evolution experiment.

The simulations also predicted which mutations would prevail. Relative to the mutant library, amplifications declined sharply and mobile-element insertions rose, in both cases to levels close to those in the evolved populations (Fig. 6B). Indels and SNPs shifted in the same direction as in evolution, though the simulations over-represented indels and under-represented SNPs. Likewise, four of the five loci most frequently mutated in the simulations—*marR* and the intergenic regions *marC/marR*, *ybjG/mdfA*, and *clpX/lon*—also carried mutations in the evolved populations (Fig. S10B). Strong antibiotic selection thus filters the available mutations in a predictable way. The one simulated locus absent from the evolved populations, *smf*, was mutated in only a single library mutant, and phenotypes estimated from a single observation are highly uncertain. Together with epistasis, this uncertainty limits the precision of the predictions. Nevertheless, mapping the local fitness landscape—the mutations available in a single step and their effects—is sufficient to predict much of how antibiotic resistance evolves.

## Discussion

By mapping the fitness and genotypes of hundreds of spontaneous resistant mutants, we found that the first step of antibiotic resistance evolution is far more diverse than previously known, yet follows simple quantitative rules. Unlike earlier approaches, this map links fitness to genotype for individual mutants across all spontaneous mutation types. The mutational target is large, dominated by amplifications and loss-of-function mutations, and includes many variants with little fitness cost (Figs. 4 and 5). Despite this diversity, a single scaling principle predicts each mutant’s fitness at any antibiotic concentration from just two parameters: its resistance (IC_50_) and its antibiotic-free growth rate (Fig. 4). Simulations built on this map, without fitted parameters, predicted the resistance gain and the number and types of mutations that fix in longer evolution experiments, where selection narrows this large target to a few high-effect routes (Fig. 6).

Covering every possible first-step mutation is not feasible, but the recurrence of mutations in the same genes suggests that we captured the most common targets of CHL resistance in *E. coli*. Notably, the genes mutated here overlap little with those identified for three other antibiotics (ciprofloxacin, nitrofurantoin, and cycloserine) in the same strain^66^, showing that the accessible mutations depend strongly on the antibiotic. First-step resistance to those drugs likely involves drug-specific targets, activation, or uptake, whereas CHL resistance is dominated by efflux. Amplifications, the most common first step in our data, went undetected in that study, because pooled sequencing dilutes the read-depth signal of an amplification carried by a single mutant. Truly comprehensive DFEs will therefore require sequencing individual mutants and creating a separate map for each antibiotic.

Our dataset also yields mutation-type-specific rates, because the relative frequencies of mutation types reflect their rates. Amplifications dominate: they arise about twice as often as SNPs and mobile-element insertions combined, and small insertions and deletions are rarer still (Fig. 5B). This supports the view that amplifications are a prominent first step in adaptation to stressors, including antibiotics^57,75^. Amplifications persist in bacterial populations at low steady-state frequencies^76^ and give rise to transient heteroresistance through resistant subpopulations with unstable copy-number increases^56,77^. Amplifications nonetheless rarely fixed in the evolved populations, and not because they were weak: their median resistance was indistinguishable from that of other mutation types (Fig. 5C). Rather, they were under-represented among the most resistant mutants, which clonal interference favors. Our simulations reproduced this depletion from the measured effects alone, without modelling their loss, indicating that being most common is not enough to fix. This largely agrees with Schenk *et al.*^78^, who found fewer structural variants in larger populations and inferred smaller effects for structural variants than for SNPs. Because their estimates came from evolved clones, already filtered by selection, they reflect the largest effects in each type; our first-step mutants, measured before selection, show that amplifications fall short only in that tail.

About two thirds of the mutations in populations evolved under sustained antibiotic selection affected genes that were also hit among our first-step mutants, suggesting that mutational paths to resistance are largely predictable. Conversely, many first-step mutations never appeared in evolved populations, likely because of clonal interference. In large bacterial populations, beneficial mutations coexist and compete, so large-effect mutations outcompete weaker ones^12,79,80^. Our simulations reproduced this loss without epistasis between mutations (Fig. S10B). Epistasis likely narrows the options further. Under diminishing returns, a mutation adds less resistance on an already resistant background^32,33,81,82^, so beneficial mutations in the ancestor may become too weak to compete. Under sign epistasis, well documented within genes though less so between them, they may even become deleterious. Predicting it would require measuring combinations of mutations. However, the first-step data alone already predict much of the phenotypic, and even mutational, trajectory under sustained drug selection.

The fitness costs of resistance mutations may also shape evolutionary trajectories. We observed the classic trade-off between resistance and antibiotic-free fitness^48^, but it was weak (Fig. 4C). Several mutants combined high resistance with low fitness costs, which is especially concerning because such mutants can persist without antibiotic pressure^83^. Because we sampled fitness–resistance space densely, the Pareto front estimates the minimal cost of resistance: only 1.7 % of antibiotic-free growth per twofold increase in resistance (Fig. 4C). Compensatory mutations can reduce this cost further^84^. Different environmental conditions and additional stressors, in contrast, can increase costs^30,85,86^. Nevertheless, our data suggest that the potential of fitness costs to counteract resistance may be over-estimated^87^.

The distribution of resistance (IC_50_) changes across mutants is largely sufficient to predict the entire DFE at antibiotic concentrations that substantially reduce growth (Fig. 4). These predictions rest on a scaling principle for DRCs. For a given antibiotic and strain, the shape of the DRC remains essentially the same in resistant mutants, and only two parameters, IC_50_ and *g*_0_, change. Our measurements confirmed this principle for hundreds of spontaneous mutants (Fig. 4), extending earlier validations for gene-deletion mutants, smaller sets of spontaneous mutants, and a small combinatorial landscape of resistance mutations in *E. coli*^37,51,55^. Once the joint distribution of IC_50_ and *g*_0_ is known, mutational effects can be predicted at any antibiotic concentration—an essential ingredient for mathematical models of evolution^37,51,88^, including under the varying concentrations that occur during therapy.

We deliberately focused on a tractable model system and an antibiotic that allow precise growth-rate measurements as a proxy for fitness. Two steps remain to bridge the gap to clinical application. First, we need to understand how epistasis between mutations reshapes the DFE. Characterizing all double mutants is infeasible, but the DFE of second mutations could be measured on a handful of backgrounds that already carry a resistance mutation. These measurements might reveal simple rules, such as global epistasis, that make measuring every combination unnecessary and sharpen predictions further. Second, the DFE and mutational targets should be mapped for clinically relevant antibiotics, pathogens, and host-like conditions. With such maps, mathematical models could forecast resistance evolution in the clinic and guide treatments that leave resistance little room to evolve^89^. Evolution is often considered inherently unpredictable. Our results suggest that, for antibiotic resistance, much of this unpredictability disappears once the local fitness landscape is mapped.

## Materials and Methods

### Strains and growth conditions

*Escherichia coli* strains are derived from BW25113^90^. Barcode library strains carry low-copy-number plasmid pCS-*λ* containing a luminescence operon (*luxCDABE*)^91^ and a DNA barcode of 16 random nucleotides interspersed with six known ones (NNNNTANNNNAC-NNNNATNNNN). Cultures were grown in LB medium or on LB agar (Sigma-Aldrich) at 30 °C. Where necessary, 25 µg ml^−1^ kanamycin was added for plasmid maintenance or selection. If not specified otherwise, strains were grown in clear, flat-bottom 96-well plates. Strains were stored as cryo stocks at −80 °C with 15 % glycerol.

### Barcode library generation

Barcodes were introduced through overhang PCR of the kanamycin resistance cassette using primers *barc-kan-fwd* (with barcode) and *kan-rev* (Table 1). The barcoded cassette was introduced into the luminescence plasmid via restriction ligation cloning. Plasmids with random barcodes were transformed into one stock of chemically competent BW25113 in six separate transformation events. Single colonies were selected and subjected to barcode validation using Sanger sequencing of colonies (LGC Genomics). Due to growth during transformation, 266 barcodes were isolated repeatedly and are represented by multiple cultures. Only cultures with high-quality Sanger sequencing results indicating unique barcodes with a Hamming distance of at least 3 to all other barcodes were used. In total, the library consisted of 617 unique barcodes.

**Table 1:**
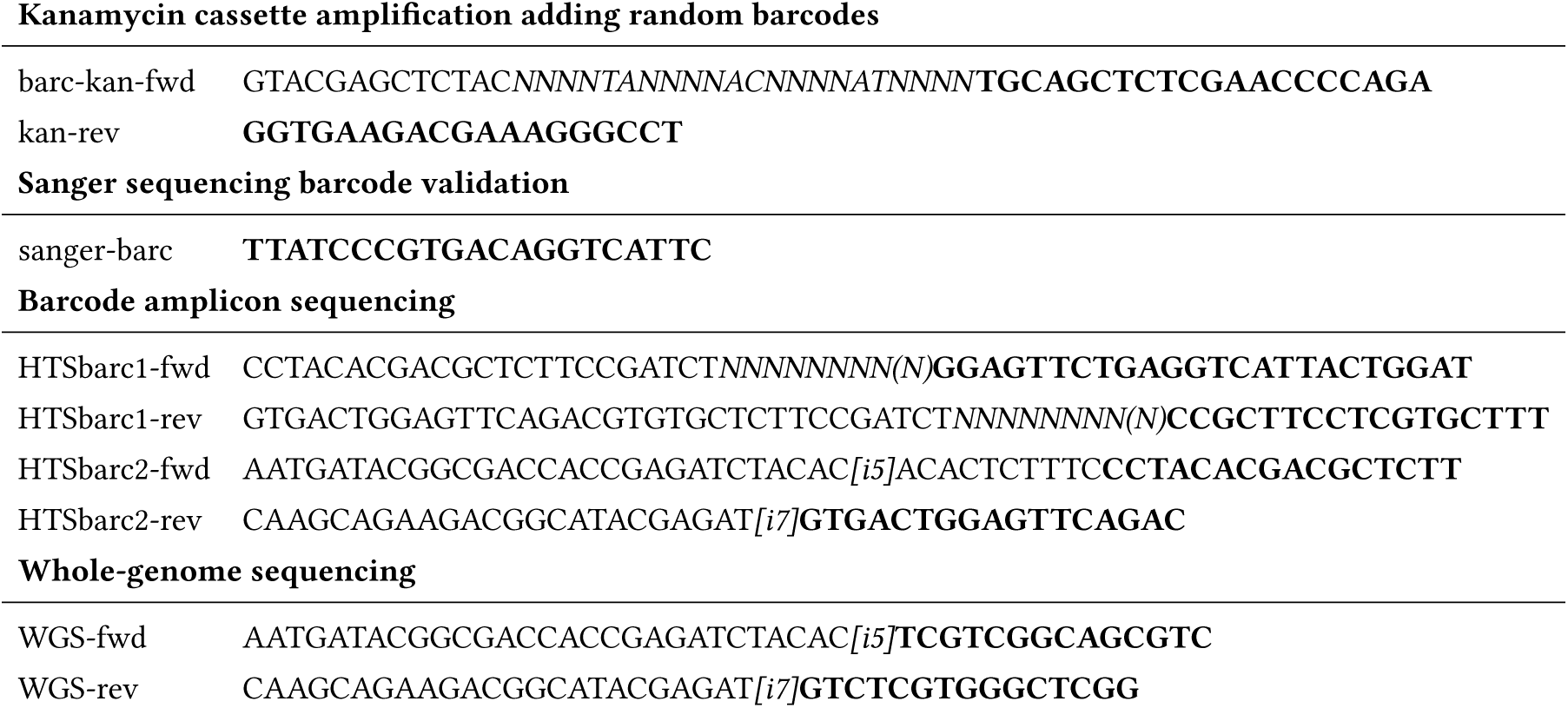
Primers used in this study. Primer sequence is listed as 5’ to 3’. The binding region is highlighted in bold and barcode, UMI, and indices in italics. Indices i5 and i7 are multiplexing indices.

### Cell number determination using Coulter Counter

Samples from growing cultures were counted using the Multisizer3 Coulter Counter (Beckman Coulter). Samples were diluted in 12 ml isotone II diluent (Beckman Coulter) to achieve counts of approximately 5 × 10^4^ to 10^5^ particles in 250 µl or 500 µl. Blank counts from isotone alone were subtracted. Note that this count includes both dead and live cells unlike plating and colony-forming-unit counting.

### Mutant selection

In six separate experiments, parallel cultures were inoculated using barcode-library cryo stocks and allowed to accumulate spontaneous mutations during growth (Fig. 1). 200 µl cultures in a 96-well plate started with approximately 1225 cells and were grown to saturation for 20 h to 22 h. Saturated cultures were diluted 1:50 or 1:25 in 384-well plates and pinned onto selection plates at the CHL wild-type MIC of 8 µg ml^−1^ and 7 µg ml^−1^ for different CHL stocks using a 1536–pin tool with a BM3-BC pinning robot (S&P Robotics Inc.). Pinning was typically repeated on two selection plates and one plate without CHL to observe contamination and confirm empty wells. This yielded 16 replicate positions per plate and culture with approximately 2000 cells per pin. After 4 d, well-separated single colonies of spontaneously resistant mutants appeared in a subset of the 1536 positions. Colonies were segmented on plate images using an intensity threshold in a custom Python script using scikit-image version 0.26.0^92^. Segments were filtered based on size and eccentricity to exclude speckles, and colony centers were mapped to the closest grid position. Colonies from independent initial cultures with different barcodes were randomly selected and automatically picked into 96-well plates for overnight growth and cryo storage. Additionally, colonies from LB plates without selection were picked as ancestor replicates and LB-only wells were used as contamination control.

### Mutation rate estimation

The beneficial mutation rate per genome per generation towards CHL resistance was determined from the high-throughput mutant selection experiments. The number of colonies per culture was counted from grid positions and image segmentation. The resulting distribution of colony numbers was compared to a simulated distribution based on the experimental procedure since it differed from the standard protocol. Instead of plating large parts of the culture and counting all colonies, small samples were taken that yielded either no or a single colony if at least one mutant cell was present. In the simulations, ancestor cells grew deterministically from an inoculum of 1225 cells for 20 generations and mutants appeared stochastically according to cell number and tested mutation rate. The mutant population expanded by a reduced, fixed, per-generation growth factor set to 1.95 to match measured fitness costs. Finally, the mutant count was scaled by the experimental dilution factor and a pin volume of 0.1 µl to give the expected number of mutants per pin. The number of pins with colonies was drawn from a binomial distribution with the probability of recovering at least one mutant in the pinned sample. The cultures of the experiment were simulated 500 times with corresponding dilution and pin sample numbers to obtain a probability mass function of colony numbers at a given mutation rate. The Jensen–Shannon distance between observed and simulated mass functions was minimized. The starting mutation rate was determined in a grid search. 95 % CIs were determined from 500 resamples of experimental cultures for mutation rate estimation. Cultures without mutant colonies were counted for the *p*_0_-method and adjusted for the fraction of cells plated over all pins divided by the total number of cell divisions to also generate an estimated mutation rate per genome per generation^14^. Confidence intervals were determined from 1000 resamples of colony numbers.

### Luminescence-based growth rate determination

Monoculture fitness was determined as exponential growth rate from luminescence based growth curves. Cultures were inoculated into white 96-well plates (Nunc) with 150 µl LB at the respective CHL concentration from pre-diluted cryo plates using a 0.2 µl pin tool for a total diluti on of 1.3 × 10^−6^. Luminescence was measured in a plate reader either in a Synergy H1 (Biotek) with continuous shaking or for multiple plates using a BioStack to Synergy Neo2 reader combination (Biotek) in a climate-controlled room at 30 °C shaking only before each luminescence measurement. Exponential growth rate was fit to luminescence curves after background subtraction using median luminescence from non-inoculated wells during the first two hours. Values were then log-transformed for a linear regression to the exponential growth phase approximately between 10^2^ to 10^5^ counts per second (cps). Luminescence measurements can detect as few as 100 to 1000 cells per well, enabling high-precision growth-rate quantification over approximately 20 generations until saturation, superior to optical density at 600 nm (OD_600_) measurements^91^.

### DNA barcode sequencing-based pooled growth rate

To determine mutant fitness, we tracked total population size and mutant frequencies at 4 to 6 time points over an average of 11 generations of exponential growth (Fig. 2). Cell numbers were determined using the Coulter counter as described above and mutant frequencies were determined based on amplicon-sequencing of the barcode.

We first validated the approach using 340 mutants and seven ancestor replicates across three drug concentrations (Fig. S1), then expanded and modified it to 474 mutants and 18 ancestor replicates to resolve high-resolution DRCs across nine drug concentrations (Fig. 2). Cryo stocks of isolated mutants and several ancestor replicates were pooled in equal volumes, spun down and re-diluted in fresh LB medium with kanamycin and glycerol. For the competition assay, the mutant pool was diluted 1:1000 to 10^6^ cells ml^−1^ in 20 ml LB at the respective CHL concentration. Cultures were grown for 9.5 h to 19.5 h, depending on the CHL concentration, with samples taken periodically. In the first competition assay, the condition without CHL was repeated twice to investigate noise from mutant growth. For 3 µg ml^−1^ and 6 µg ml^−1^ CHL, the volume of larger samples was replaced with fresh LB at the appropriate concentration. In the second competition assay, the three lowest concentrations were diluted 1:10 with fresh medium at the respective CHL concentration before reaching stationary phase, to increase the number of generations and thereby improve the resolution of the small fitness differences in these conditions. For high concentrations, sampling was restricted to fewer generations, before a few mutants came to dominate the population. All main figures show data from the second experiment.

Amplicons for high-throughput sequencing (HTS) were generated in two PCR steps. Heated culture samples were directly used as templates in the first PCR. For this, samples were spun down, resuspended in 20 µl sterile water and heated at 99 °C for 10 min. 15 µl or less, depending on cell density, was added to a standard 30 µl Q5 polymerase reaction (New England Biolabs) with primers HTSbarc1-fwd and HTSbarc1-rev for two PCR cycles. These amplify the barcode region adding parts of the Illumina adaptors and unique molecular identifiers (UMIs) to uniquely label each template (Table 1). We used primers with UMIs lengths of 8 or 9 to create a read offset and increase template diversity during amplicon sequencing^93^. PCR products were then purified using magnetic beads (Ampliclean, Nimagen or NucleoMag® NGS Clean-up and Size Select, Macherey-Nagel). In the second PCR amplification, the Illumina adaptor was completed with variable index combinations for multiplexing. 7.5 µl to 16 µl of the purified amplicon were used in a 25 µl Q5 PCR for 23 cycles. Samples were gel purified to completely remove primer-primer amplicons using spin-columns (Macherey-Nagel). PCR reactions were sensitive to cell numbers, and several had to be repeated to achieve sufficient product. Amplicons were sequenced on an Illumina MiSeq v2, 2 x 150 bp paired-end reads at the Cologne Center for Genomics (CCG).

Fitness was determined by fitting exponential growth rates to mutant or ancestor population sizes. Paired-end reads were merged using NGmerge (0.3)^94^. The region around the barcode was identified in the amplicon reads using regular expression matching allowing up to three mismatches. Extracted barcodes were then matched to the mutant library. We removed reads from PCR amplification by counting UMIs with up to one nucleotide difference only once. Cell number counts with re-dilution were translated into a single longer growth curve and used to calculate mutant population size with mutant barcode frequencies. The exponential growth phase was then fit using a log-linear regression to estimate fitness over concentrations (LinearRegression from Python package scikit-learn 1.9.0). The standard error of this regression fit is reported. Eight mutants with few reads or repeated barcodes identified during whole-genome sequencing (WGS) were excluded from further analysis.

### Dose–response curve measurement

CHL dependent growth rates either from barcode assays or luminescence measurement were fit with a Hill function of the form *g*(*c*) = *g*_0_*/* [1 + (*c/*IC_50_)*^n^*] to generate DRCs. Here, *g*_0_ is the antibiotic-free growth rate, *n* the Hill exponent, *c* the antibiotic concentration, and IC_50_ the half-maximal inhibitory concentration. The package lmfit (version 1.3.4)^95^ was used with *g*_0_ fit constrained to two standard deviations of ancestor replicate measurements around the strain-specific measured antibiotic-free growth rate. Low-cost mutations were analyzed by linear regression of the mutants on the Pareto front, which was determined with a maxima-of-a-point-set algorithm. To compare a more similar inhibition level between ancestors and mutants (Fig. S3), we repeated the Hill function fit excluding fitness values below 0.2 normalized to antibiotic-free growth rate of the respective strain.

### Whole-genome sequencing

WGS was performed to identify resistance mutations. DNA was extracted from 50 µl overnight culture of mutants. Cells were pelleted in U-bottom elution plates and lysed in 25 µl lysis buffer with final concentrations of 80 % PBS (Sigma-Aldrich), 4 mM Tris-HCl pH 8 (Calbiochem), 0.4 mM EDTA (Carl Roth), 0.3 % Triton-X 100 (Sigma-Aldrich), 5 mg ml^−1^ lysozyme (Sigma-Aldrich) and 0.5 mg ml^−1^ proteinase K (Macherey-Nagel). Cells were incubated at 37 °C for one hour with occasional shaking and at 56 °C for at least six hours or overnight. Genomic DNA was extracted using components and protocol of the kit NucleoMag® DNA Bacteria (Macherey-Nagel) in a down-scaled version of the protocol using 13.5 µl of the lysed sample. DNA was eluted in 15 µl IME and processed following the Hackflex v1 protocol^96^ with volumes further reduced by half. In brief, genomic DNA was tagmented using BLT beads from Nextera® DNA Flex Library Prep (Illumina), amplified adding combinatorial dual indexes using PrimeSTAR GXL DNA Polymerase (Takara) and individually cleaned with size-selection using SPRIselect beads (Beckman Coulter). Finally, 5 µl of each sample were pooled and sequenced on a NovaSeq X+ 10B lane with 150-bp paired-end reads (NGSelect Ready-to-Load, Eurofins Genomics).

WGS data was filtered to exclude low-quality and adapter sequences using fastp^97^ (version 0.24.0, default settings except for enabled adapter detection for paired end reads, minimum read length 40 bp, poly-G trimming, cutting by quality score from front and back). We identified mutations and structural variations using breseq^98,99^ (version 0.39.0). Reads were annotated to a deposited genome of BW25113 (NCBI accession: CP009273), after adding IS elements (ISEscan 1.7.3)^100^, and the plasmid pCS-*λ*, constructed according to the cloning procedure above. In a first round, mutations present in the ancestor chromosome and plasmid were identified and added to the reference files. Breseq was run in polymorphism mode to detect mutations. Read alignment was limited to a coverage of 300 to reduce run-time. Due to the pooling strategy reads per sample varied considerably. Samples that did not yield a coverage above 30 were excluded from further analysis (20 % of samples). To remove mutations from sequencing errors while considering incompletely fixed mutations, we considered all genomic mutations with a frequency of at least 30 %. 34 mutations identified on the plasmid were not considered further, given their neutral signature with a high number of synonymous mutations. Junctions in the *fimE*/*fimA* switch as well as a prophage repetitive region stfP/stfE were excluded even if their frequencies were higher since they were also present at low frequencies in ancestor replicates. Additional junctions were manually inspected and curated into mutations. Knockout and promoter predictions from breseq were used directly.

We predicted amplifications based on coverage, breseq junctions and AmpliFinder 1.0.0^101^ in a custom script. We identified regions with at least 1.3-fold coverage compared to the median. Coverage was smoothed using the function median_filter from scipy.ndimage with size 5000 (scipy version 1.18.0). Close regions of increased coverage were merged. Precise start and end positions were identified using amplifinder results, breseq junctions or IS elements in the genome. Detected amplifications and positions were manually curated and added to the list of mutations from breseq. In total, 344 mutations were detected, comprising 120 unique mutations (Table S1).

In 137 mutants, no mutation was identified. Given that CHL is bacteriostatic for this strain, ancestor cells may survive the mutant-selection procedure and could expand in frequency during regrowth of the colony for storage and sequencing. Together, colonies underwent approximately 40 generations, during which costly mutations would be selected against. However, the IC_50_ of these clones differed significantly from that of both the ancestor replicates and the mutants with detected mutations (Figs. 5C and S5). This indicates that, while some clones may retain the ancestral genotype, many such clones carry undetected mutations. In particular, short-read sequencing often misses structural rearrangements, especially in repetitive DNA sequences^99^. Furthermore, resistance mutations may revert during the CHL-free growth periods after selection. Notably, amplifications frequently had copy numbers below two indicating a certain rate of reversal as observed previously^56^. Thus, many of these cases may come from loss of amplifications below the detection limit.

Mutations were compared to data from Lukačišinová *et al.*^33^. They sequenced 56 samples from populations evolved to adaptively increasing CHL concentration. We considered only 47 samples from 180 h of evolution and reanalyzed the uploaded data (PRJEB37495) following the procedure described above. Sequenced samples contain 20 different gene-deletion strains with 2 to 3 replicates and 7 replicates of their reference strain Δ*lacA* BW25113, which is expected to behave like our ancestor. We pooled data from all strains assuming that most gene deletions do not change mutational effects. Mutations in the evolution experiment were matched to the mutant library by considering first identical mutations, then inactivating mutations and finally mutations in the same gene or genomic region for amplifications. If the mutant library contained multiple matching mutations, their effects were averaged. IC_50_ fold changes of matched mutations were combined multiplicatively, weighted by their frequency in the evolved population, to predict an expected resistance increase for each sequenced population.

### Statistical analysis

Correlations were analyzed using Pearson’s *r* or Spearman’s rank correlation (*r*_s_) from scipy.stats (1.18.0) depending on whether a linear relationship was expected. When comparing predictions for the same data or reproducibility of repeated measurements we determined concordance correlation (*r*_c_)^102^. In all cases, permutation tests with 9999 resamples were used to calculate *p*-values, yielding a minimum *p*-value of 2 × 10^−4^. Correlation between amplicon fitness effect and amplicon size was determined using Spearman’s *r*_s_ on median values for repeated amplifications weighted by √n according to the number of occurrences of each amplification. Here an exact permutation test was performed across all possible resamples.

The tail of the DFE was analyzed using the likelihood-ratio test for deviation from exponential following Beisel *et al.*^46^. Exponential and generalized Pareto distributions were fit using scipy.stats fixing loc to 0 by shifting the distribution according to the chosen threshold. LRT p-values come from 1999 resamples adjusted for multiple testing with Holm’s method yielding a floor of 0.0045. Shape values fit across thresholds were compared to assess variability. 95 % CI error bars around shape parameter values were determined from 500 resamples.

Significant differences between two groups, i.e., ancestor and mutant Hill exponents as well as repeatedly mutated genes or genes selected during the evolution experiment were determined using a two-sided Mann–Whitney *U* test (scipy.stats 1.18.0) accounting for multiple testing where appropriate using Holm’s method (multipletests from statsmodels 0.14.6). Mutation number and types were compared using a Kruskal–Wallis test on *g*_0_ and IC_50_, Holm-adjusted for multiple testing. Dunn’s test (scikit-posthocs 0.14.0), also Holm-adjusted, was used as a post hoc test to identify differences between groups.

The relationship between mutation frequency and phenotype was tested with a linear regression (scipy.stats) and *p*-value from Pearson’s *r* as described above. The relationship between copy number and phenotype was tested with an ordinary least-squares model (ols from statsmodels 0.14.6) with a common slope for copy number and a separate intercept for each amplicon (ANCOVA). *p*-value comes from these results. In both cases, log-transformed IC_50_ values were used.

### Stochastic simulations of evolution experiments

Evolutionary trajectories were simulated following the experimental procedure of the evolution experiment. Cells grew in batch cultures between median OD_600_ values of 0.0096 to 0.1160 leading to a dilution rate of 1:12 and approximately 3.6 generations per growth phase. The simulations were limited to 130 generations, matching the generations after 180 h of evolution. Cell numbers in experimental cultures of 200 µl were estimated from OD_600_ calibration reported in Stevenson *et al.*^103^ for mid-log phase cells adjusting for a Tecan plate reader. This yields a population size fluctuating between 1.5 × 10^6^ to 1.8 × 10^7^. The simulations started from an ancestral population that grew stochastically in small time steps. Dividing cells could mutate at the estimated mutation rate (Fig. 1). Each mutation created a new lineage whose fitness effect and IC_50_ fold change were multiplied with those of any previous mutations. Lineages then continued to grow and mutate stochastically, accumulating multiple mutations. Mutations were drawn from the mutant library, using the mean effect for repeated mutations and accounting for their higher frequency. After each growth phase, populations were diluted by randomly removing cells from lineages and the concentration was updated according to the current population growth rate to maintain 50 % growth inhibition. The ancestor Hill exponent was used throughout. Reported results are from 1000 simulation runs. The example Muller plot was created with PyFish 1.2^104^.

## Acknowledgements

We thank Joachim Krug and Francesca Macaluso for feedback on the manuscript; and the entire Bollenbach and Krug groups for fruitful discussions. We thank Yuval Mulla for support with the WGS procedure. This work was funded by the Deutsche Forschungs-gemeinschaft (DFG, German Research Foundation - SFB1310/3 – 325931972). We further-more thank the ITCC (IT Center University of Cologne) for providing compute resources on the DFG-funded HPC (High Performance Computing) system RAMSES (Research Accelerator for Modeling and Simulation with Enhanced Security) as well as support (DFG funding number: INST 216/512-1 FUGG).

## Author contributions

Conceptualization: T.F. and T.B. Investigation: T.F. Formal Analysis: T.F. Method development: T.F. and J.A. Visualization, data curation and software: T.F. Writing – original draft: T.F. Writing – review and editing: T.F., J.A., and T.B. Supervision: T.B. Funding acquisition: T.B.

## Supplements

### Supplementary Tables

**Table S1: Mutations identified by whole-genome sequencing.** Each row lists one mutation in one strain. Strains with several mutations span several rows, and strains without a detected mutation appear once with mutation type none. Point mutations, small insertions and deletions, MOB insertions and large deletions were called with breseq; amplifications were called separately (Methods). genomic_mutnum: number of mutations detected in the strain; copy_number: estimated copy number of the amplified region; frequency: fraction of reads supporting a breseq call. Provided as Supplementary_Table_S1.xlsx.

**Table S2: Dose–response parameters and fitness estimates.** Each row is one strain. Columns g_0, IC50 and exponent give the fitted Hill parameters, each with the standard error of the fit (suffix _se). Columns g_0_normed and IC50_foldchange give *g*_0_ and IC_50_ relative to the ancestor mean. Columns c_loglin_fitness give the fitness at chloramphenicol concentration *c* (in µg ml^−1^), estimated by log-linear regression (Methods), and columns c_ll_fit_stderror give its standard error. Columns labeled exp1 come from the first competition experiment with two drug-free replicates (0A, 0B) and two concentrations. Genotypes are summarized as mut_type:gene_name (AMP:start-end for amplifications), with multiple mutations separated by semicolons; none marks sequenced strains without a detected mutation, and not-sequenced marks strains without whole-genome data. Provided as Supplementary_Table_S2.xlsx.

### Supplementary Figures

**Figure S1:**
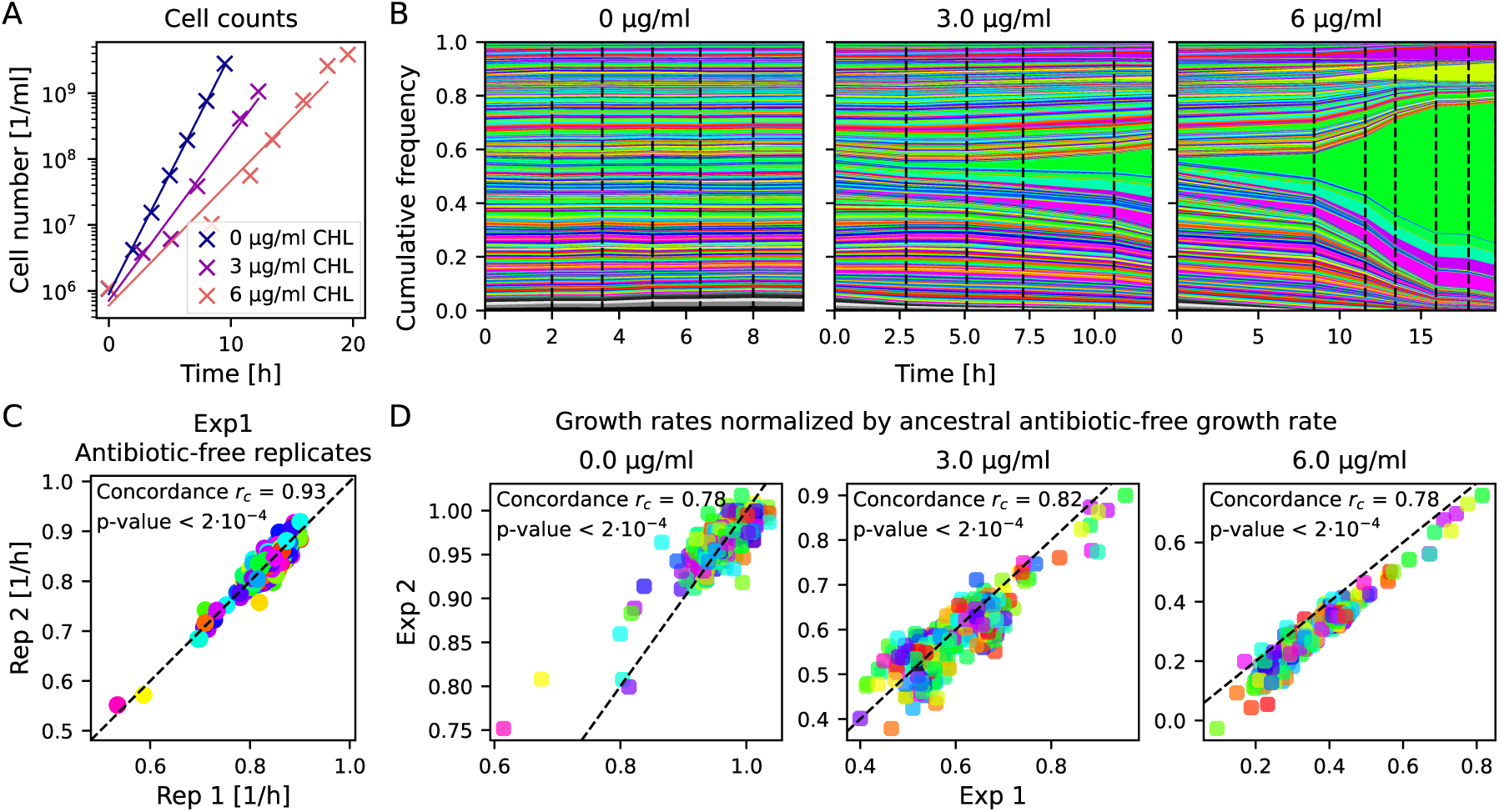
The pooled barcode competition assay is highly reproducible. **A–C)** First barcode experiment. **A)** Total cell numbers used to estimate mutant population sizes. Without antibiotic, cells were not counted at the sequencing time points, so cell numbers were interpolated by log-linear regression. **B)** Cumulative mutant frequency over time at each antibiotic concentration. Dashed vertical lines mark sampling time points; gray at the bottom, ancestor replicates. **C)** Fitness of the same barcoded library in two independent antibiotic-free cultures (340 mutants, 7 ancestor replicates). Agreement is higher than between separate experiments (D). **D)** Fitness in the first and second barcode experiments at the three concentrations measured in both (0 µg ml^−1^, 3 µg ml^−1^ and 6 µg ml^−1^; 336 mutants, 3 ancestor replicates). Growth rates were generally higher in the second experiment, owing to less frequent sampling and better temperature control, so all growth rates were normalized to the antibiotic-free growth rate of the ancestor in the same experiment (0.87 h^−1^ and 1.13 h^−1^, respectively). Concordance correlation coefficients (*r*_c_) are given in each panel; significance was assessed by a permutation test.

**Figure S2:**
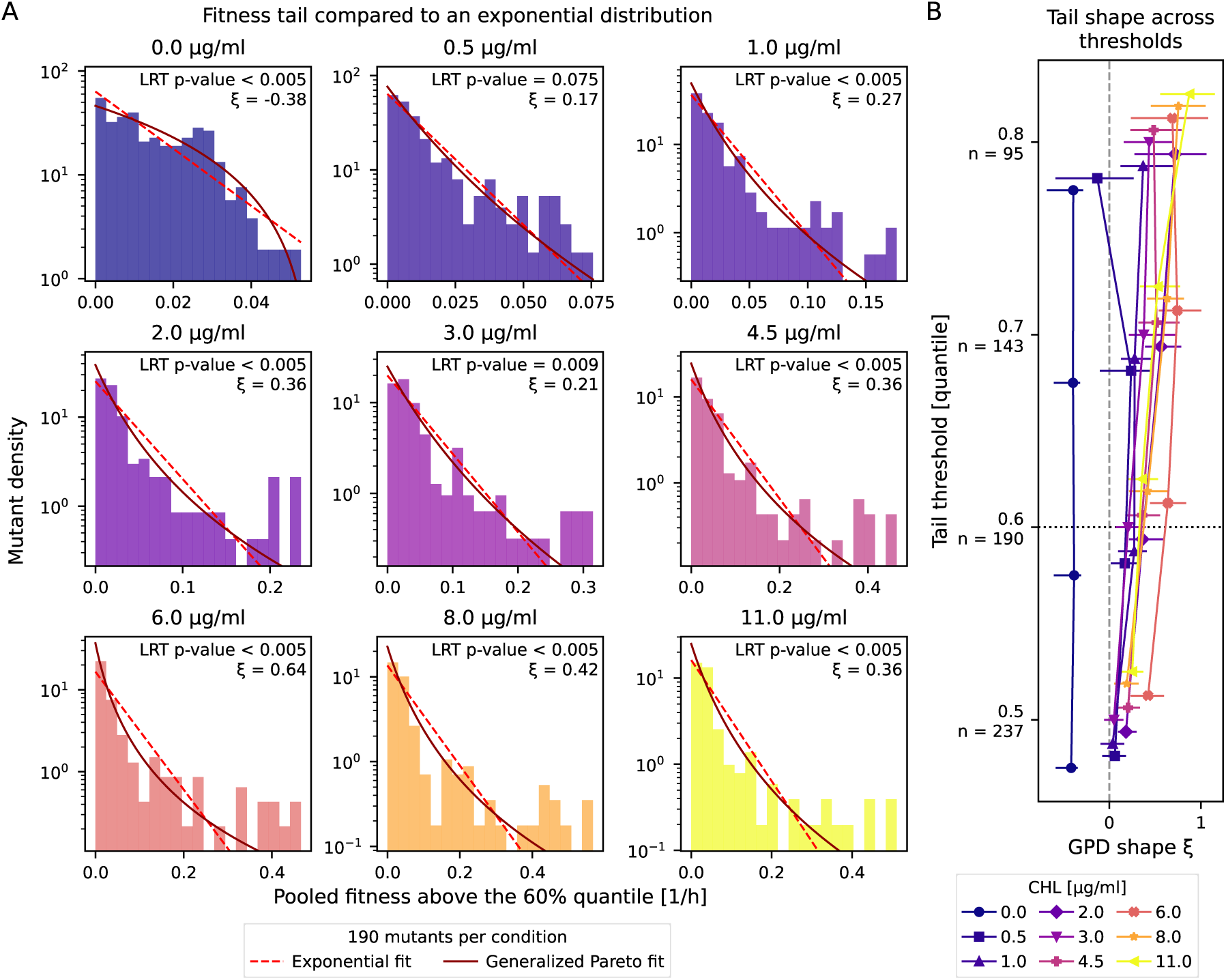
The beneficial tail of the DFE becomes heavy under antibiotic stress. **A)** Beneficial tails of the pooled fitness distributions at all measured concentrations, comprising mutants with fitness above the 60th percentile. Each tail was fit with an exponential and a generalized Pareto distribution, and the two fits were compared by a likelihood-ratio test. *ξ* is the shape parameter of the generalized Pareto fit: *ξ <* 0 indicates a bounded tail, *ξ* = 0 an exponential tail, and *ξ >* 0 a heavy tail. **B)** Shape parameter *ξ* for different thresholds defining the tail. Error bars are 95% CIs from bootstrapping.

**Figure S3:**
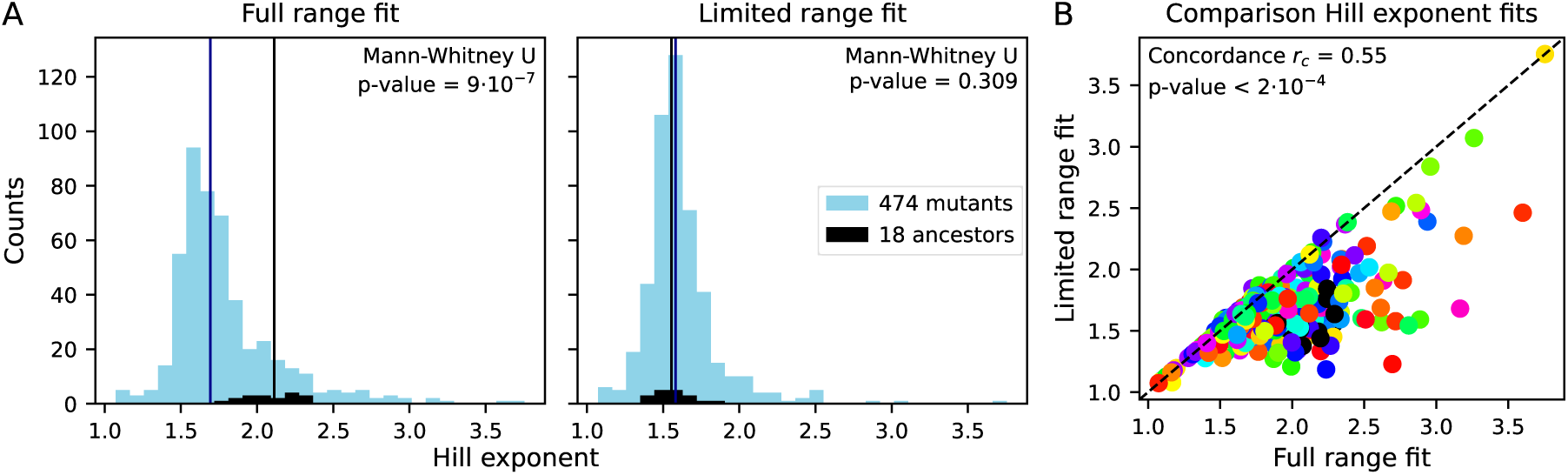
Mutant Hill exponents match the ancestor’s when fits are restricted to comparable inhibition. **A)** Hill exponents from fits to the full DRCs or only to growth rates inhibited by less than 80 %. With the restricted range, mutant and ancestor distributions do not differ significantly. **B)** Hill exponents from the two fits; *r*_c_ is given in the panel. For many mutants, restricting the range barely changes the exponent.

**Figure S4:**
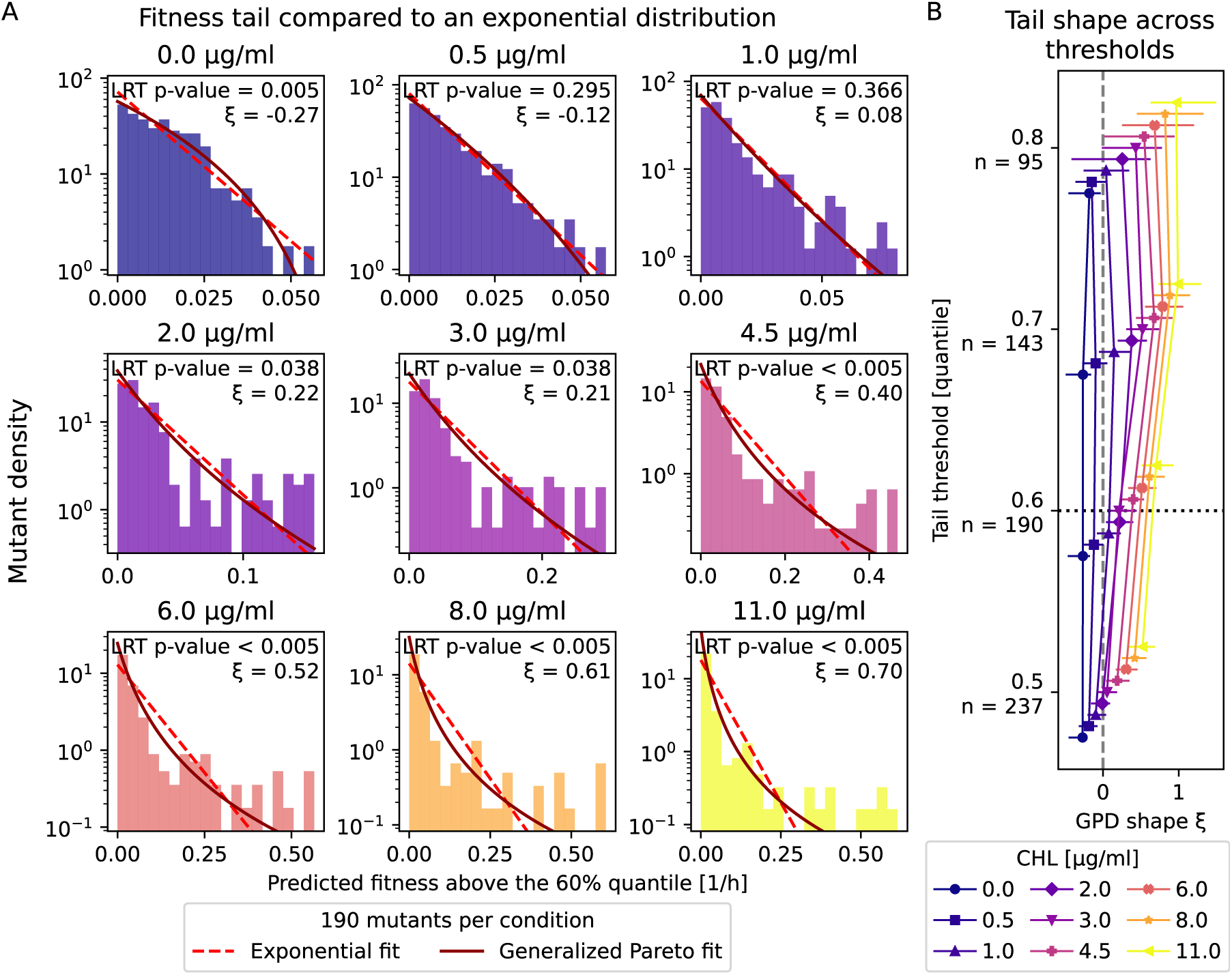
The DRC scaling principle reproduces the shift from a bounded to a heavy DFE tail. **A)** Beneficial tails of the predicted fitness distributions at all measured concentrations, comprising the 190 mutants with predicted fitness above the 60th percentile. Fitness was predicted for each mutant from its own *g*_0_ and IC_50_ together with the ancestor’s Hill exponent (Fig. 4). Each tail was fit with an exponential and a generalized Pareto distribution, and the two fits were compared by a likelihood-ratio test. *ξ* is the shape parameter of the generalized Pareto fit: *ξ <* 0 indicates a bounded tail, *ξ* = 0 an exponential tail, and *ξ >* 0 a heavy tail. **B)** Shape parameter *ξ* for different thresholds defining the tail, with the number of mutants in the tail given at each labeled threshold. Dashed vertical line, *ξ* = 0; dotted horizontal line, the 60th percentile used in A. Error bars are 95 % CIs from bootstrapping. Across concentrations the predicted *ξ* rises from −0.27 to 0.70, following the measured trend (Fig. S2); the predicted tails are significantly bounded without drug and significantly heavy from 2 µg ml^−1^ upwards.

**Figure S5:**
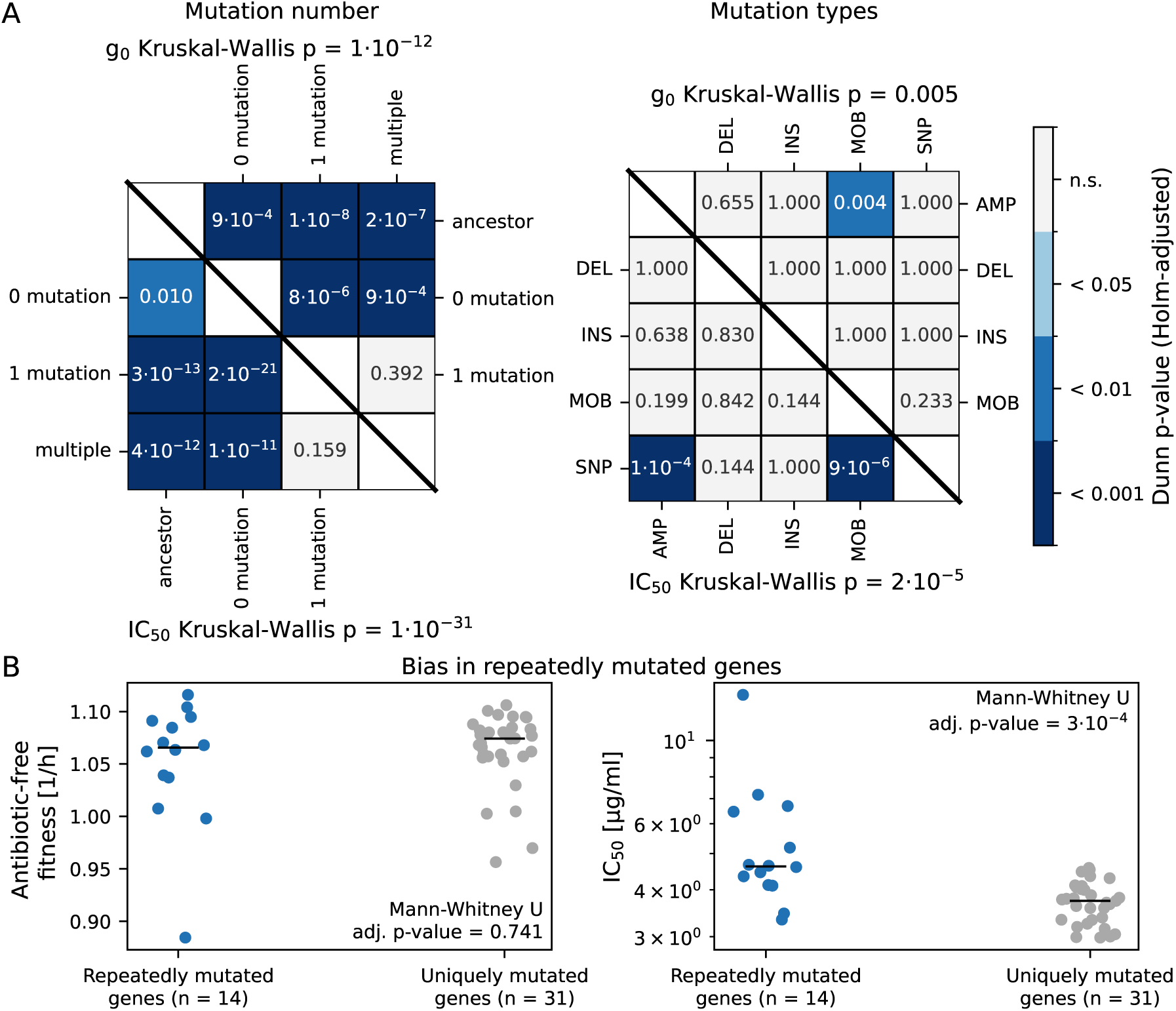
Statistical comparison of mutant fitness and resistance. **A)** Antibiotic-free fitness (*g*_0_, upper triangle) and resistance (IC_50_, lower triangle) of the ancestor and of mutants grouped by number of detected mutations (left) and by mutation type (right). Kruskal–Wallis tests detected significant differences in every comparison, and pairwise differences were then identified with Dunn’s post hoc test. *p*-values were corrected for multiple testing with Holm’s method. **B)** *g*_0_ and IC_50_ for genes mutated once or repeatedly in the library considering only mutants with a single mutation; for repeatedly mutated genes, the mean per gene is used.

**Figure S6:**
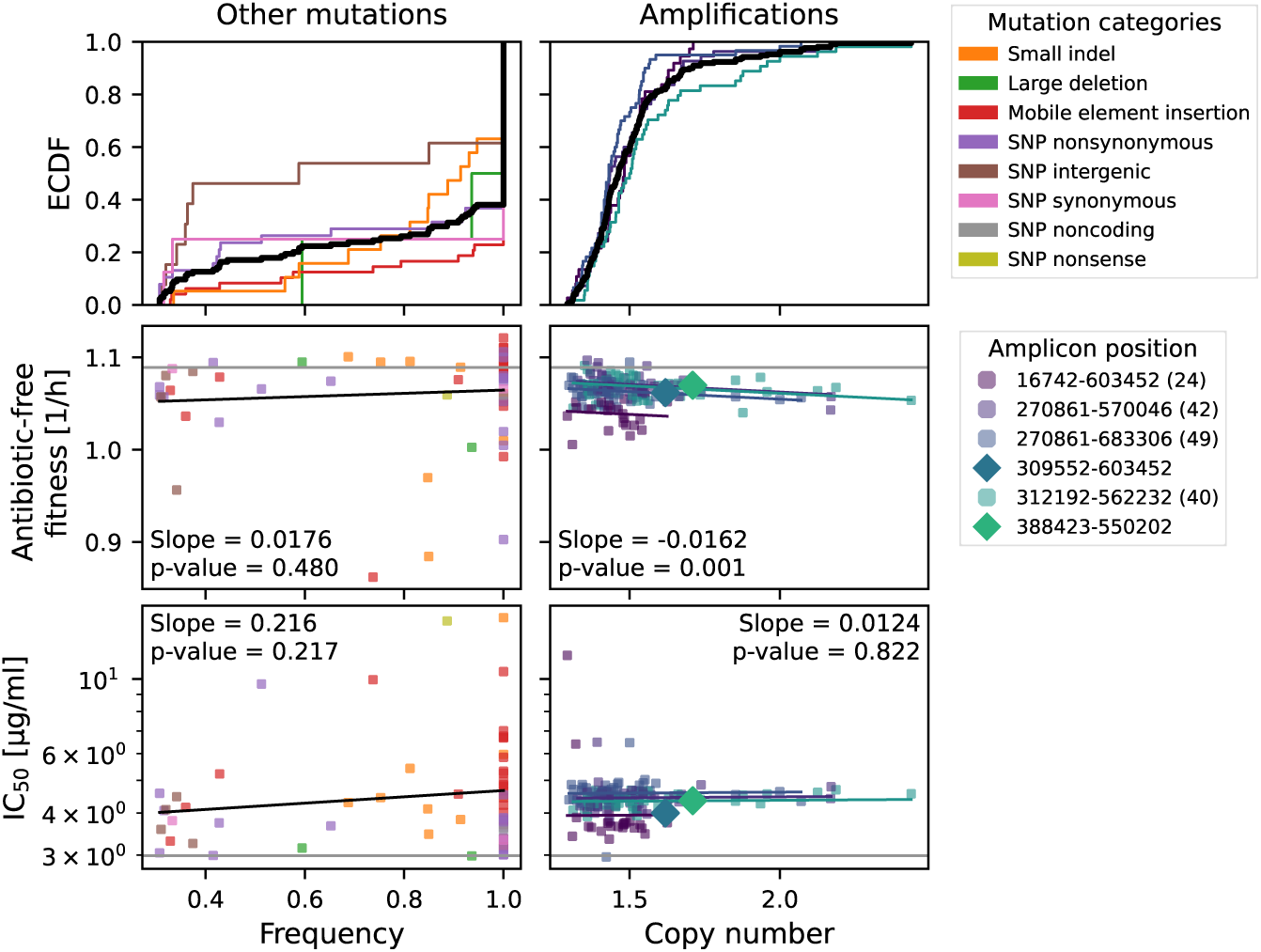
Mutation frequency does not predict mutant phenotypes, whereas higher amplicon copy number slightly lowers fitness. Left, all mutation categories except amplifications, against mutation frequency; right, amplifications, against amplicon copy number. Top, cumulative distributions over all identified mutations (frequency, *n* = 134; copy number, *n* = 210): most mutations are at or near frequency one, and most copy numbers are below two. Colored lines, individual mutation categories or amplicons represented by more than three mutants; black, all pooled. Middle and bottom, antibiotic-free fitness (*g*_0_) and resistance (IC_50_) of phenotyped mutants carrying a single mutation; gray lines, ancestor median; circles, recurrent amplicons; diamonds, amplicons found in a single mutant. Black lines, linear regression across all non-amplification mutations, with *p*-values from a permutation test on Pearson’s correlation; colored lines, analysis of covariance across the amplifications with a common slope and separate intercepts per amplicon. IC_50_ is plotted on a logarithmic axis and was log-transformed for regression. Fitness and resistance were independent of mutation frequency, as was resistance of copy number; only fitness declined with increasing copy number, slightly but significantly (slopes and *p*-values in each panel).

**Figure S7:**
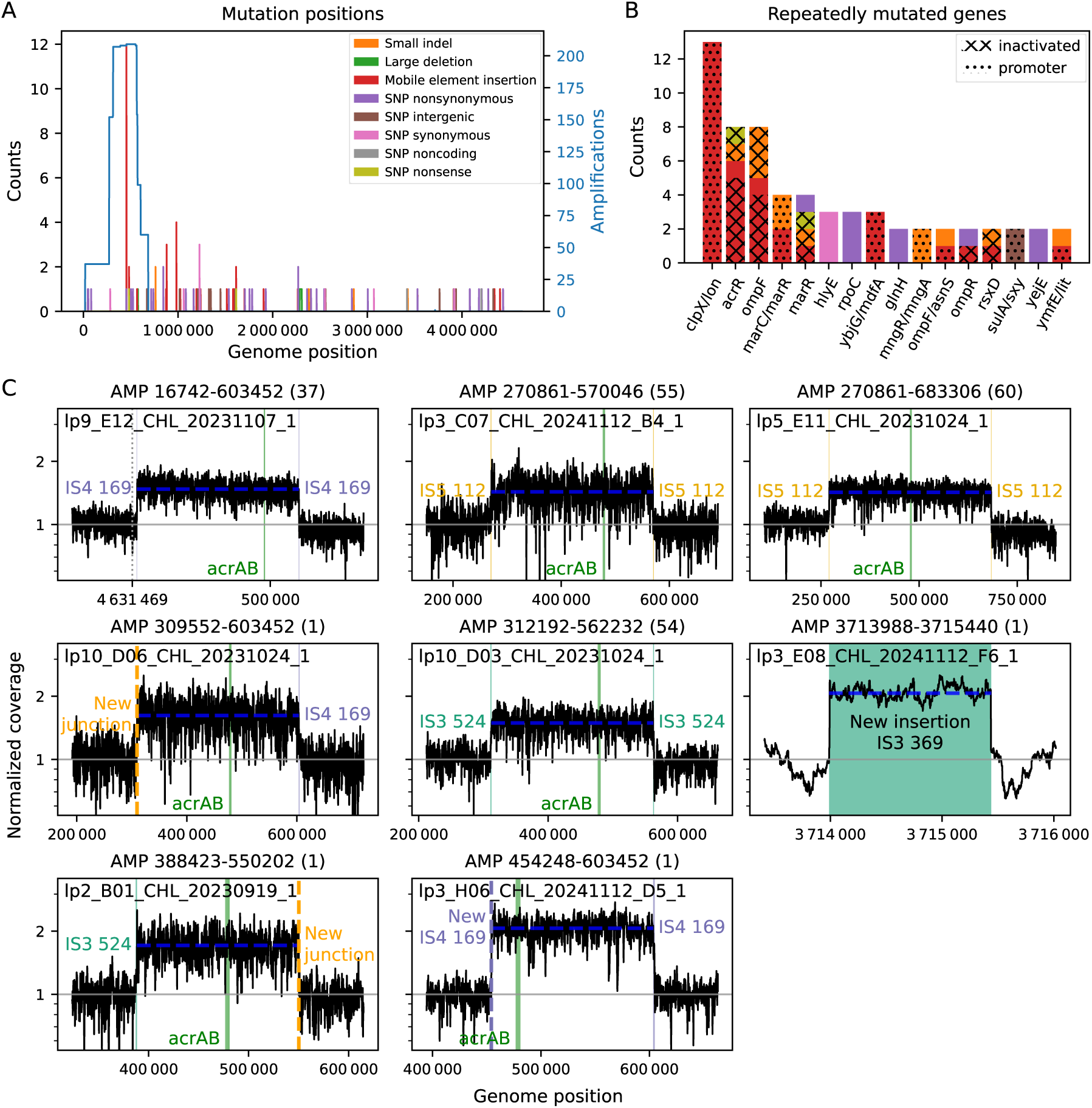
Amplifications in the mutant library cluster around *acrAB*. **A)** Genomic positions of all detected mutations. Other mutations are spread across the genome, whereas amplifications cluster in one region (counts on right axis). **B)** Number of mutants for each repeatedly mutated gene or region, inactivation and promoter mutations as predicted by breseq. **C)** Read coverage, normalized to the genome median, for each of the eight distinct amplifications, with the number of mutants carrying it. Each plot shows a representative mutant with intermediate copy number (mutant ID, top left); dashed blue lines mark the median copy number of the amplicon. Dotted gray line, connection of circular genome. Recurrent amplicons are bounded by pre-existing IS elements, whereas single amplicons are bounded by new junctions or newly inserted IS elements. Seven of the eight amplicons, accounting for 209 of 210 amplification events, include *acrAB*, and most raise its copy number less than twofold (median 1.46).

**Figure S8:**
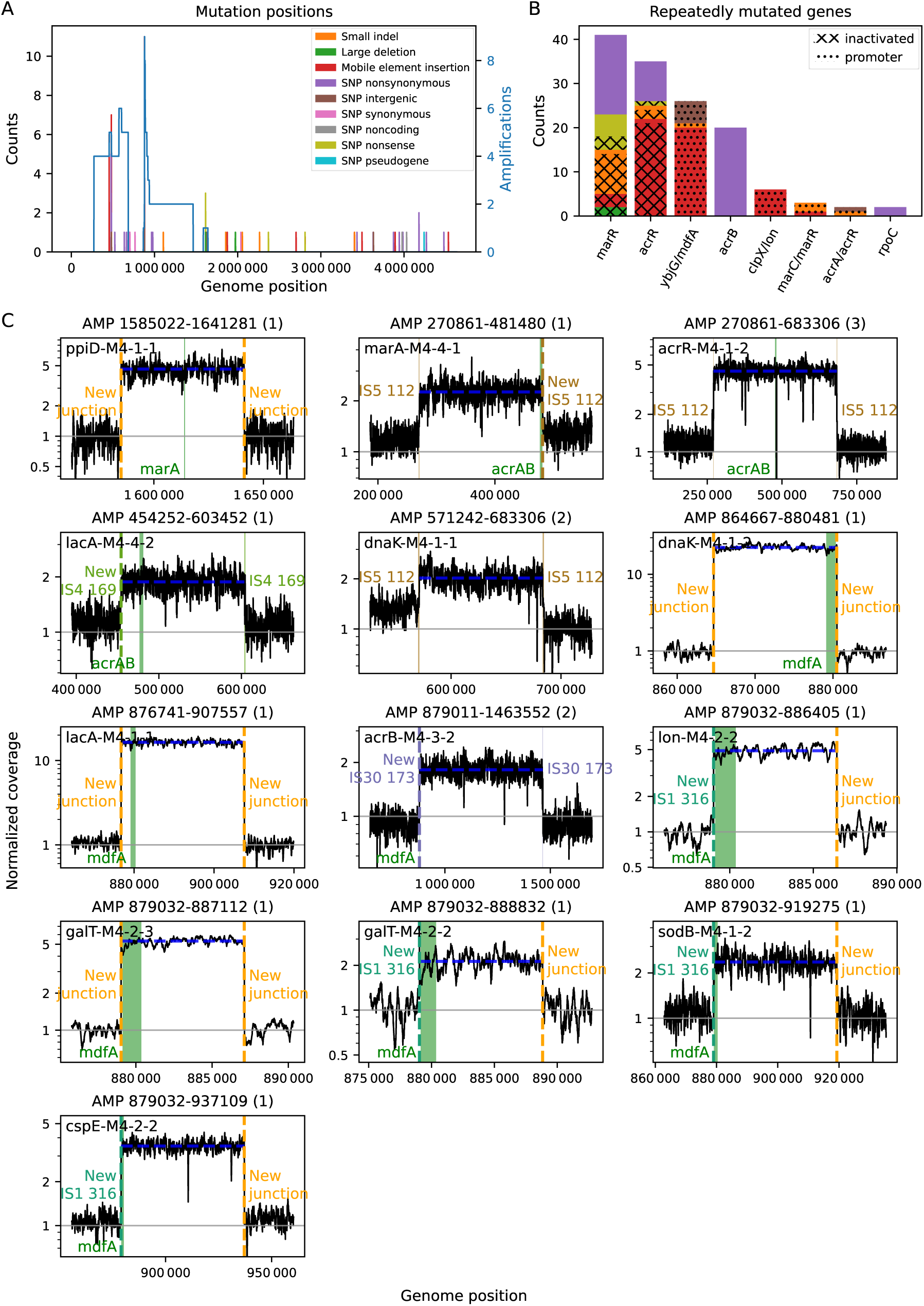
Amplifications in the evolved populations of Lukačišinová *et al.* ^33^. **A)** Genomic positions of all detected mutations. Mutations are less widely spread than in the mutant library, and amplifications are often shorter and cluster in two regions, around *acrAB* and *mdfA* (counts on right axis). **B)** Number of populations for each repeatedly mutated gene or region, inactivation and promoter mutations as predicted by breseq. **C)** Read coverage, normalized to the genome median, for each distinct amplification, with the number of populations carrying it. Each plot shows a representative population with intermediate copy number (sample ID, top left); dashed blue lines mark the median copy number of the amplicon. Amplicon borders are formed by IS elements or new junctions. Label top left, evolution experiment sample shown^33^.

**Figure S9:**
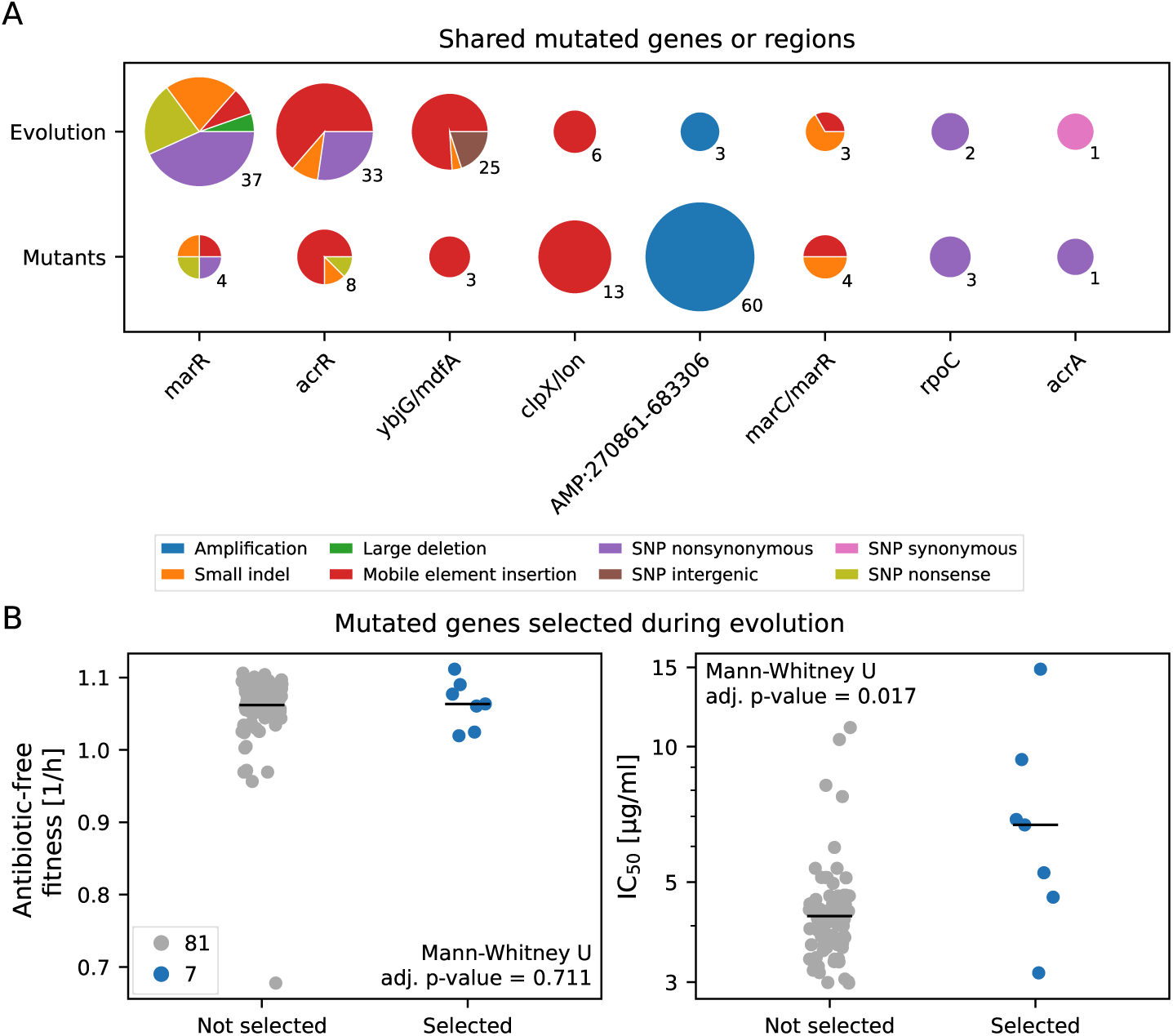
Genes mutated in both the mutant library and the evolved populations. **A)** Number and types for mutations shared on the level of genes or region in both the library and the evolved populations. Pie size reflects the number of repetitions with precise value in lower right corner of each pie. **B)** Antibiotic-free fitness (*g*_0_) and resistance (IC_50_) of library mutants in genes or regions that were or were not selected in the evolved populations; for genes mutated more than once, the mean is used. For *acrA* no phenotypic information was determined.

**Figure S10:**
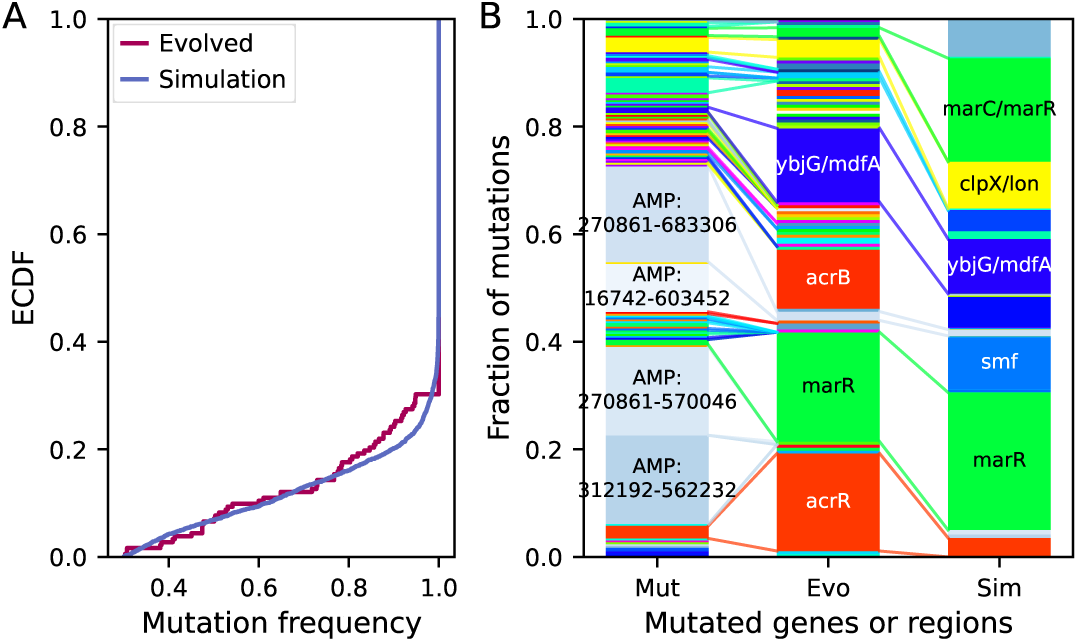
Simulations reproduce the mutation frequencies and gene targets of the evolved populations. **A)** Cumulative distributions of mutation frequency in the evolved populations (*n* = 185) and in the simulations (*n* = 3137 from 1000 runs). In both, most mutations are at or near fixation, and the rest form a tail of intermediate-frequency mutations carried by subpopulations. The 30 % frequency threshold of the sequencing analysis was applied to the simulations as well. **B)** Mutated genes or regions in the mutant library (Mut), the evolved populations (Evo), and the simulations (Sim). Each segment is one gene or region, scaled by its share of all mutations in that group; lines connect the same gene or region across groups. Mutations in the library are spread across many loci and dominated by amplifications, whereas the evolved and simulated populations each concentrate on a few.

